# Pharmaceutical agent cetylpyridinium chloride inhibits immune mast cell function by interfering with calcium mobilization

**DOI:** 10.1101/2023.05.23.541979

**Authors:** Bright Obeng, Christian M. Potts, Bailey E. West, John E. Burnell, Patrick J. Fleming, Juyoung K. Shim, Marissa S. Kinney, Emily L. Ledue, Suraj Sangroula, Alan Y. Baez Vazquez, Julie A. Gosse

**Affiliations:** Department of Molecular and Biomedical Sciences, University of Maine, Orono, ME, USA; Department of Biology, University of Maine Augusta, Augusta, ME, USA

**Keywords:** Antimicrobial, calcium, cetylpyridinium chloride, quaternary ammonium compound, degranulation, endoplasmic reticulum, inositol 1,4,5-triphosphate, cytoplasmic pH, mast cell, microtubule, plasma membrane potential.

## Abstract

Cetylpyridinium chloride (CPC) is an antimicrobial used in numerous personal care and janitorial products and food for human consumption at millimolar concentrations. Minimal information exists on the eukaryotic toxicology of CPC. We have investigated the effects of CPC on signal transduction of the immune cell type mast cells. Here, we show that CPC inhibits the mast cell function degranulation with antigen dose-dependence and at non-cytotoxic doses ∼1000-fold lower than concentrations in consumer products. Previously we showed that CPC disrupts phosphatidylinositol 4,5-bisphosphate, a signaling lipid critical for store-operated Ca^2+^ entry (SOCE), which mediates degranulation. Our results indicate that CPC inhibits antigen-stimulated SOCE: CPC restricts Ca^2+^ efflux from endoplasmic reticulum, reduces Ca^2+^ uptake into mitochondria, and dampens Ca^2+^ flow through plasma membrane channels. While inhibition of Ca^2+^ channel function can be caused by alteration of plasma membrane potential (PMP) and cytosolic pH, CPC does not affect PMP or pH. Inhibition of SOCE is known to depress microtubule polymerization, and here we show that CPC indeed dose-dependently shuts down formation of microtubule tracks. *In vitro* data reveal that CPC inhibition of microtubules is not due to direct CPC interference with tubulin. In summary, CPC is a signaling toxicant that targets Ca^2+^ mobilization.

## 1. Introduction

Despite widespread use (American Chemical Society, 2021) of antimicrobial cetylpyridinium chloride (CPC) at millimolar or higher concentrations (Rawlinson et al., 2008), the eukaryotic toxicology of this drug is largely unstudied.

CPC is used as an antimicrobial at millimolar levels in dozens of personal care products applied directly to humans (including mouthwash, toothpaste, lozenges, nasal sprays, cleansing wipes, ointments, lip balm, hair products, deodorant) (Bernauer et al., 2015; Mao et al., 2020; Pubchem, 2023). Human foods such as poultry are bathed in CPC up to 23 millimolar for disinfection purposes (Food and Drug Administration, 2023), and custodial staff work directly with high-CPC cleaning solutions. Despite this widespread use, a lack of proven CPC antibacterial efficacy compared to plain cleansers led the FDA to effectively ban CPC from hand soap in 2016 (Wolf, 2016) and hand sanitizers in 2019 (Gottlieb, 2019).

CPC does exhibit specific beneficial uses in humans in particular as an anti-gingivitis (Teng et al., 2016) and an anti-halitosis agent (Yaegaki & Sanada, 1992). CPC has also exhibited anti-viral efficacy, against influenza in mice (Popkin et al., 2017) and zebrafish (Raut et al., 2022) and against SARS-CoV2. CPC has been investigated in human clinical trials as a mouthwash treatment to reduce SARS-CoV-2, with mixed results: CPC did not affect salivary viral load but did significantly reduce viral infectivity at 1-hour post-mouthwash (Sánchez Barrueco et al.) and elevated saliva levels of nucleocapsid protein (which indicates disruption of viral particles) (Alemany et al., 2022).

Thus, while there are known or potential benefits of CPC, the risks are unknown due to the lack of published eukaryotic toxicological data. The benefits of any widespread, high-dose chemical must be weighed with potential risks by gathering data, including on underlying biochemical mechanisms.

CPC, a quaternary ammonium salt, contains a positively charged nitrogen in its headgroup and a long hydrocarbon tail (Pubchem, 2023). CPC acts as a detergent (Mao et al., 2020) when above its critical micelle concentration (CMC) of 600-900 µM (Abezgauz et al., 2010; Mandal, 1991; Mukerjee, 1971; Shi et al., 2011), thereby killing bacteria. However, the CPC effects on mast cells described in this article are not caused by the detergent action of CPC because experiments in this study were conducted with CPC doses (0.1-10 µM) far lower than the CMC. Furthermore, CPC doses used in this study are not cytotoxic to various cell types, as determined by trypan blue exclusion (Raut et al., 2022), lactate dehydrogenase release (Raut et al., 2022), ToxGlo (Weller et al., 2022), and plasma membrane potential (Figure 5B) assays. Thus, the CPC exposures used in this study are affecting signal transduction, not cell viability.

CPC is retained in human tissue: following brief (minutes) mouthwash application, low- µM levels of CPC were slowly released into saliva, out of the oral mucosa of human volunteers, over 24 hours (Bonesvoll & Gjermo, 1978). Also, CPC administered orally to mice at a relevant level (1 mg/kg) resulted in low-µM CPC levels in the blood within hours (Pottel et al., 2020). Based upon this mouse pharmacokinetic study and levels of CPC in chicken for consumption, we estimated that a typical American may have a blood CPC concentration ranging from ∼0.1 μM - 0.3 μM due to chicken consumption alone (Weller et al., 2022). CPC administered at sub-µM levels to zebrafish via swim water led to behavioral changes and altered brain neurotransmitter levels (Dong et al., 2022). Thus, despite a lack of direct human exposure and pharmacokinetics studies, emerging evidence suggests that CPC undergoes uptake and distribution throughout the body. Moreover, related quaternary ammonium compounds have been detected in human blood at nM levels, which were associated with biochemical effects in humans (Hrubec et al., 2021).

Despite widespread CPC usage/ingestion and likely uptake into body tissues and fluids, human epidemiology studies of CPC are currently lacking (PubMed,2023). Also, there exist very few mechanistic studies of CPC on eukaryotes. However, CPC does cause pulmonary toxicity in rats (Kim et al., 2021; Lin et al., 1991) and modulates human gingival fibroblast prostaglandin production a function also of mast cells (Kim et al., 2005).

We previously showed that CPC inhibits functions of mast cells (MCs) (Raut et al., 2022), which are ubiquitous: MCs are found in lung, skin, gastrointestinal mucosa, neurological, and other tissues (Blank & Benhamou, 2013; Kuby, 1997; Theoharides et al., 2012). Thus, MCs are poised for CPC exposure via inhalation, food ingestion, and product application. Mast cells are involved in immune and neurological responses and diseases (Krystel-Whittemore et al., 2015; Metcalfe et al., 1997; Theoharides et al., 2016). These cells possess surface receptors, FcεRI, which, when activated, trigger signaling, leading to the release of stored bioactive mediators within cytoplasmic granules via exocytosis, a process known as degranulation. These mediators include serotonin, histamine, and β-hexosaminidase (Abramson & Pecht, 2007).

Degranulation (illustrated in Figure 7) of mast cells begins when FcεRI (Kuby, 1997; Stevens & Austen, 1989) receptors bind IgE (Keown et al., 1998) and then are crosslinked by multivalent antigen (Ag; allergen) (Ravetch & Kinet, 1991), kicking off a tyrosine phosphorylation cascade which activates phospholipase C gamma 1 (PLCγ1). PLCγ1 binds phosphatidylinositol 4,5-bisphosphate (PIP_2_) in the inner leaflet of the plasma membrane (PM) and catalyzes its hydrolysis, generating inositol-1,4,5-trisphosphate (IP_3_) (Gibson et al., 1994). Crucially, IP_3_ binds its receptors on the endoplasmic reticulum (ER) (Scharenberg et al., 2007), activating their Ca^2+^ channel function such that Ca^2+^ flows from the ER into the cytosol (Berridge, 1993). The reduced ER [Ca^2+^] induces conformational change of stromal interaction molecule 1 (STIM1), which then binds the Orai1 subunit of the Ca^2+^ release-activated Ca^2+^ (CRAC) channel (Putney, 1986), opening it to a flood of external Ca^2+^ into the cytosol (Hogan et al., 2010; Vig et al., 2006). This process, “store-operated Ca^2+^ entry,” or SOCE (Clapham, 1995; Putney, 1986, 1990), is required for degranulation (Holowka et al., 2012). Additionally, mitochondrial Ca^2+^ uptake, first from ER Ca^2+^ efflux and secondly via SOCE (Takekawa et al., 2012), through the mitochondrial Ca^2+^ uniporter (MCU) (Furuno et al., 2015), enhances degranulation(Furuno et al., 2015; Takekawa et al., 2012). Ca^2+^ mobilization is a key event both for MC degranulation and for various functions of most eukaryotic cells (Kozak, 2018).

A key portion of the driving force for SOCE is the plasma membrane potential (PMP), or the voltage difference across the PM (Nelson, 2017). Dissipation of the established PMP in mast cells inhibits cytosolic Ca^2+^ rise and, thus, degranulation (Mohr & Fewtrell, 1987). Cytosolic pH level is also crucial to activity of the CRAC channel (Beck et al., 2014): acidification of the cytosol inhibits Ag-stimulated cytosolic Ca^2+^ rise in MCs (Sangroula et al., 2020).

Polymerized microtubules serve as the “railroad tracks” along which granules are transported to the PM for exocytosis (Smith et al., 2003). Ag-stimulated increase in cytosolic [Ca^2+^] is required to stimulate binding of positive regulator protein Git1 to tubulin, consequently causing enhanced polymerization and, thus, degranulation (Martin-Verdeaux et al., 2003; Smith et al., 2003; Sulimenko et al., 2015; Tasaka et al., 1991; Urata & Siraganian, 1985).

We previously showed that positively-charged CPC disrupts the interactions of the negatively-charged lipid PIP_2_ with three of its protein binding partners (myristoylated alanine-rich C-kinase substrate, hemagglutinin, a construct containing a pleckstrin homology domain) (Raut et al., 2022). Thus, in this investigation*, we hypothesized that CPC electrostatically interferes with PIP_2_-dependent SOCE*, as a biochemical explanation for CPC effects on degranulation. As other alternative explanations for CPC inhibition of degranulation, we have investigated CPC effects on PMP, pH, and microtubules.

In this study, the RBL-2H3 (rat basophilic leukemia cells, clone 2H3; “RBL”) mast cell model is employed because these cells are functionally homologous to and contain the core functional machinery of mature human MCs, rodent mucosal MCs, and of basophils (Abramson & Pecht, 2007; Metcalfe et al., 1997; Metzger et al., 1982; Seldin et al., 1985). RBL cells and primary bone-marrow derived mast cells (BMMCs) respond similarly to environmental stimuli (Alsaleh et al., 2016; Thrasher et al., 2013; Zaitsu et al., 2007). Thus, RBLs are scientifically suitable for toxicological/ pharmacological studies of mast cell signal transduction.

CPC inhibits immune cell function (Raut et al., 2022); however, the biochemical mechanisms underlying CPC’s inhibition of signal transduction are not yet known. Determination of the underlying biochemical mechanisms of CPC can lead to predictions of its impact on other cell types. In this study, we defined conditions (Ag and CPC doses, timing effects) under which CPC modulates mast cell degranulation. We tested the hypothesis that CPC disruption of PIP_2_ or PMP or cytosolic pH leads to impaired Ca^2+^ mobilization and Ca^2+^- dependent events, including microtubule polymerization, essential for mast cell degranulation. Here, we examined the effects of CPC in mast cells on Ca^2+^ mobilization and dynamics in the ER, mitochondria, and cytosol. We aimed to discover CPC effects on SOCE via multiple novel techniques, including developing a new plate reader-based mitochondrial-Ca^2+^ assay and using genetically-encoded voltage indicators (GEVI), novel in this field (Society of Toxicology, 2023). Our work demonstrates that CPC impairs mast cell degranulation and Ca^2+^ dynamics at doses as low as 3000-fold lower than the dosages currently administered via personal care products.

## 2. Materials and Methods

### 2.1. Cetylpyridinium chloride preparation

Cetylpyridinium chloride (CPC; 99% purity, VWR; CAS no. 123-03-5) was prepared in aqueous Tyrodes buffer (Hutchinson et al., 2011) on each experimental day, as described (Raut et al., 2022). CPC concentrations were verified for accuracy with UV-Visible spectroscopy (Raut et al., 2022). Bovine serum albumin (BSA) was added to diluted CPC solutions, resulting in BSA-Tyrodes (BT) CPC containing solutions.

### 2.2. Cell culture

RBL-2H3 mast cells were cultured as detailed (Hutchinson et al., 2011). Phenol red-free media (swapping a phenol red-free minimum essential media, MEM, for regular MEM, as utilized in (Kennedy et al., 2012)) is utilized as noted in subsequent sections to reduce background noise in microscopy experiments.

All assays that involved in-cell measurements employed fluorescent protein reporter constructs with β-barrel structures that protect the fluorophore. These constructs, based upon GFP and related proteins, are generally resistant to external perturbants (Brejc et al., 1997; Chalfie & Kain, 2005; Weatherly et al., 2018; Weatherly et al., 2016) and, thus, are robust for studies of drug or toxicant effects.

### 2.3. Degranulation assay

RBL-2H3 mast cell degranulation responses were measured as previously described in (Weatherly et al., 2013), adapted and validated for use with CPC (Raut et al., 2022). Some samples, denoted “P&C,” in graphs (Figure 1A and 1B), had CPC pretreatment for 30 min prior to the 1 h Ag/CPC exposure; “C” means CPC exposure only during the hour of Ag.

**Figure 1.**
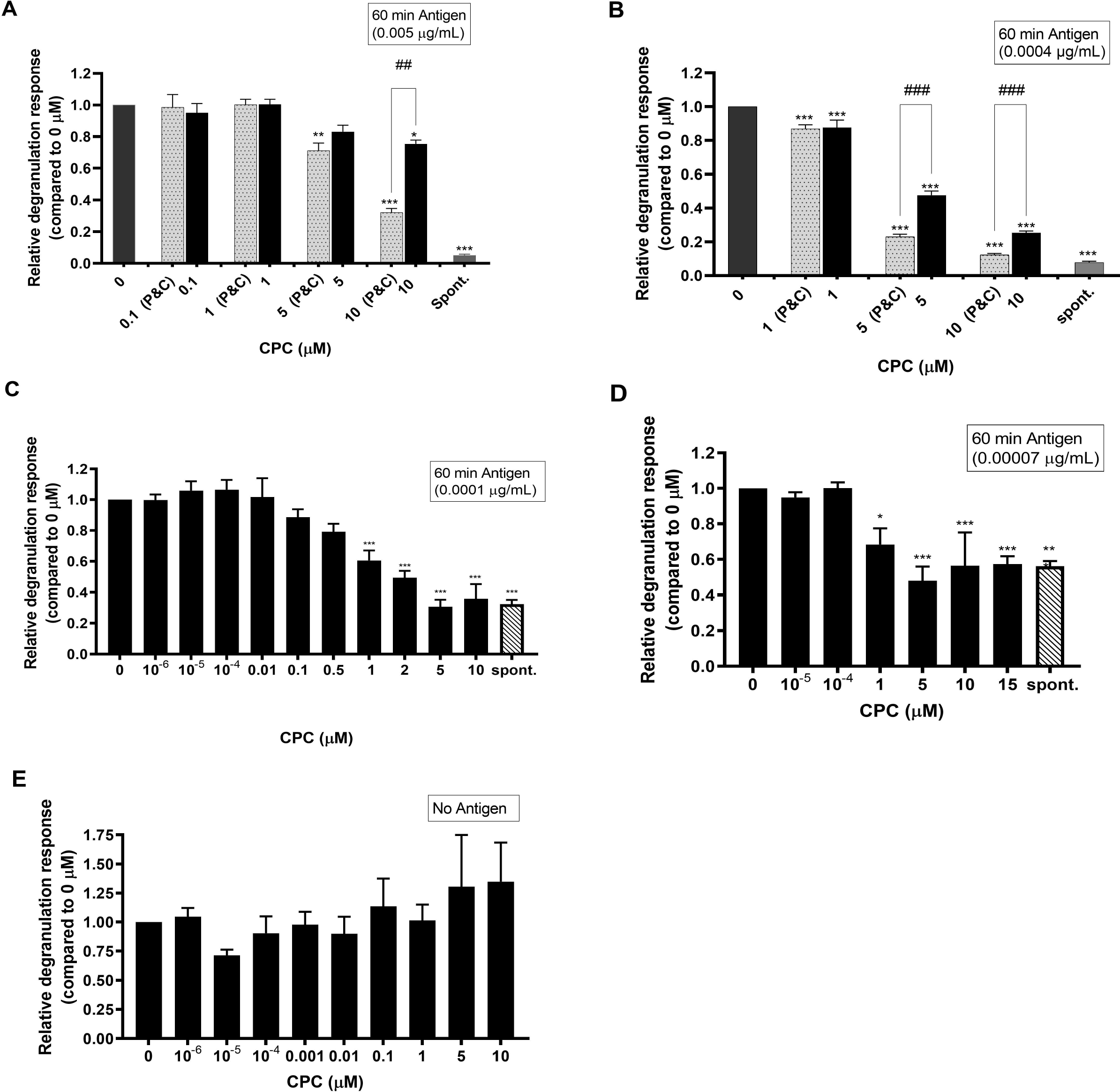
Relative degranulation response of RBL mast cells exposed to different non-cytotoxic micromolar doses of CPC. Cells were sensitized with IgE for 1 h and stimulated with either 0.005 µg/mL Ag for 1 h **(A)**, 0.0004 µg/mL Ag for 1 h **(B)**, 0.0001 µg/mL Ag for 1 h **(C)**, (or 0.00007 µg/mL Ag for 1 h **(D)**. For spontaneous release (spont. on x-axes), non-IgE-sensitized cells were exposed to CPC for 1 h in the absence of Ag **(E)**; spontaneous release measurements also appear for comparison **(A-D)**. Some samples, denoted “P&C”, in graphs **(A)** and **(B)**, had CPC pretreatment for 30 min prior to the 1 h Ag/CPC exposure. Values presented are means normalized to control, ± SEM, of at least 3 independent days of experiments; each experiment was performed in triplicate. Statistically significant results compared to the 0 µM control of each graph are represented by *p < 0.05, **p < 0.01, and ***p < 0.001 or compared to samples indicated by vertical lines are represented by ^##^p < 0.01 and ^###^p < 0.001, each determined by one-way ANOVA followed by Tukey’s post-hoc test.

### 2.4. Endoplasmic Reticulum Ca^2+^ assay

Endoplasmic reticulum (ER) Ca^2+^ levels were monitored as previously detailed (Weatherly et al., 2018). RBL-2H3 mast cells were transiently transfected with the ER-targeted genetically encoded Ca^2+^ indicator ER-GCaMP6–210 (a generous gift from Dr. Jaime de Juan-Sanz and Dr. Timothy A. Ryan) (de Juan-Sanz et al., 2017), using RBL-specific Amaxa Nucleofector transfection kit T (Lonza). In this experiment, before the 1 h ± CPC/Ag exposure and fluorescence measurement, cells were pre-treated with CPC or BT for 30 minutes. Data analysis was performed as described (Weatherly et al., 2018).

### 2.6. Mitochondrial Ca^2+^ assay

To measure mast cell mitochondrial Ca^2+^ levels in response to Ag and CPC, we developed a novel plate reader-based assay. RBL-2H3 cells were transfected with the mitochondrial calcium indicator pCMV CEPIA2mt (a gift from Masamitsu Iino; Addgene plasmid # 58218; http://n2t.net/addgene: 58218; PRID:Addgene_58218) (Suzuki et al., 2014) via nucleofection using Amaxa Nucleofector kit T (Lonza), as done in (Weatherly et al., 2018). “Mock” transfected cells underwent the same electroporation process but without plasmid DNA. Following transfection, cells were plated into 96 black-walled, clear bottom, tissue culture-treated plates (Grenier Bio-One) in 200 µL phenol red-free media at 100,000 cells/well and incubated overnight at 37 °C/5% CO_2_.

Following the overnight incubation, cells were washed twice with fresh BT, then sensitized with anti-DNP mouse IgE (Sigma Aldrich) for 1 h. After the IgE sensitization, cells were washed with BT, and pretreated with CPC or BT for 30 minutes at 37 °C/5% CO_2_. After the 30 min, the pre-treatment solutions were removed, and cells were exposed to 200 µL BT ± CPC/Ag then immediately placed into the plate reader (Synergy 2, Biotek) for fluorescence measurement. Note that “Mock” transfected cells were treated with BT alone for this hour. Excitation of 485 ± 20 nm and emission of 528 ± 20 nm were used to measure the fluorescence at 42 sec intervals over 1 h. Average background fluorescence from “Mock” transfected cells (which accounts for background fluorescence from the cells, the plate, and the BT) was subtracted from all samples at all time points before plotting (Figure 3A). To calculate mitochondrial Ca^2+^ level via area under the curve (AUC, Figure 3B) in GraphPad Prism, the y-axis baseline was defined as the average fluorescence from unstimulated cell samples.

### 2.7. Cytosolic Ca^2+^ assay

Cytosolic Ca^2+^ levels were measured as previously described (Weatherly et al., 2018). RBL-2H3 cells were transiently transfected with the genetically encoded Ca^2+^ indicator pGP-CMV-GCaMP6f (a gift from Douglas Kim & GENIE Project; Addgene plasmid # 40755; http://n2t.net/addgene:40755; RRID:Addgene_40755), (Chen et al., 2013), using RBL-specific Amaxa Nucleofector transfection kit T (Lonza). Some samples, denoted “P&C,” in graphs (Figure 4A and 4B), had CPC pretreatment for 30 min prior to the 1 h Ag/CPC exposure; “C” means CPC exposure only during the hour of Ag. Average background fluorescence from untransfected cells was subtracted from all time points. To calculate cytosolic Ca^2+^ level via area under the curve (AUC, Figure 4B) in GraphPad Prism, the y-axis baseline was defined as the average fluorescence from unstimulated cell samples.

### 2.8. Confocal microscopy

For the genetically-encoded voltage indicator (GEVI) and microtubule polymerization assays, an Olympus FV-1000 confocal microscope with Olympus IX-81 microscope and a 30-milliwatt multi-argon laser was used to collect images of transiently transfected RBL-2H3 mast cells. Transfected cells were imaged using oil immersion 100x objective with N.A 1.4, 488 nm excitation, and 505-605 nm band pass emission filter. An ibidi plate heating system maintained the temperature at 37°C during image acquisition.

### 2.9. Plasma membrane potential and cytosolic pH assay

The effect of CPC on plasma membrane potential and cytosolic pH were determined in tandem using the pH-sensitive genetically-encoded voltage indicator (GEVI) ArcLight-A242 (a gift from Vincent Pieribone; Addgene plasmid # 36857; http://www.addgene.org/36857) (Jin et al., 2012) as detailed (Sangroula et al., 2020). ArcLight fluorescence was measured for 30 min of CPC treatment. Image analysis was performed as detailed (Sangroula et al., 2020).

### 2.10. Confocal of microtubule polymerization

Microtubule polymerization was assessed as previously described (Weatherly et al., 2018), employing the construct pIRESneo-EGFP-alpha Tubulin (a gift from Patricia Wadsworth; Addgene plasmid #12298; https://www.addgene.org/12298/) (Rusan et al., 2001). Cells were plated in 8-well ibidi plates at 133,333 cells per well in 200 µL of phenol red-free RBL media, then grown overnight at 37 °C, 5% CO_2_. In this experiment, before imaging, cells were incubated with Ag ± 5 or 10 μM CPC solution in BT for 1 h at 37 °C, while unstimulated cells were incubated in plain BT. Following the 1 h incubation, cell images were collected within 20 min. Cells undergoing microtubule polymerization were determined by looking at both fluorescent and DIC images and using blinded manual classification by two separate researchers. At least 38 cells for each treatment group were analyzed.

### 2.11. Tubulin polymerization assay

*In vitro* tubulin polymerization assays were performed according to the manufacturer’s instructions (Cytoskeleton, Inc.) optimized for polymerization inhibitors, and similarly to previously published methods (Weatherly et al., 2018). The final assay conditions in calcium-free buffer were pH 6.9, 1.1 mM GTP, 1.58 mg/ml tubulin protein, 18.4% glycerol, 1.8 mM 4-(2-hydroxyethyl)-1-piperazineethanesulfonic acid (HEPES), 72 mM Piperazine-N,N’-bis[2-ethanesulfonic acid] sesquisodium salt (PIPES); 0.5 mM ethylene glycol-bis(b-amino-ethyl ether) N,N,N’,N’-tetra-acetic acid (EGTA), 2.0 mM magnesium chloride (MgCl_2_), and 5.9 μM of the proprietary fluorophore, mixed together before plating in a black-walled 96-well plate (Corning Costar). An enhancer of tubulin polymerization, paclitaxel, was used as a positive control. Wells also contained final concentrations of 0 μM CPC (control), 5 μM CPC, or 2.7 μM paclitaxel–which were added to assay plate wells first as concentrated (11X), 5 μL volumes, followed by 50 μL volumes of the master tubulin mix noted above. Apart from CPC, all assay chemicals were provided within the kit, other than HEPES (BioBuffer Solutions). Final total volumes per well were 55 μL. Fluorescence intensity was measured every 42 seconds for a duration of 1 hour at 37°C with a plate reader (Synergy 2, Biotek) (Temperature 37°C, Excitation 360/40 nm, Emission 460/40 nm, Gain 60, Shake every 5 seconds, Optics position Top 50%).

Data observed in duplicate were averaged and then background-subtracted using the 0 μM CPC time point zero. Background is time point 0 min of 0 μM CPC group. All data points were normalized to the 0 μM CPC final time point (60 min), then data from 4 independent experiments were used to plot normalized fluorescence intensity against time (Figure 6E). Duplicate-averaged, background-subtracted RFU values of the first 30 min of measurement were used to calculate the rate of tubulin polymerization (Figure 6F). This polymerization rate was compared between the 0 μM CPC group and the 5 μM CPC group using a two-tailed t-test for difference between means (Figure 6F). Duplicate-averaged, background-subtracted RFU values of the last 10 min of measurement were utilized to represent total tubulin polymerization. Total tubulin polymerization was compared between the 0 μM CPC group and the 5 μM CPC group using a two-tailed t-test for difference between means (Figure 6G).

### 2.12. Statistical analyses

All analyses were performed in Graphpad Prism as detailed (Weatherly et al., 2018). Biological replicates from at least three different experimental days were averaged and used to determine SEM. As indicated in individual figure legends, significance levels were assessed via t-tests or one-way ANOVA with Tukey’s post hoc tests, as appropriate. At least 33 cells per treatment type underwent image analysis in Figures 5 and 6.

## 3. Results

### 3.1. CPC inhibits degranulation in RBL mast cells in an exposure time- and antigen dose-dependent manner

Non-cytotoxic concentrations of CPC dose-dependently inhibit degranulation in RBL cells stimulated by a moderate antigen dose, 0.0004 µg/mL, within an hour of co-exposure (Raut et al., 2022). In this study, we further examined the effects of CPC on mast cell degranulation upon stimulation by various antigen doses and exposure timeframes, using our published fluorescence-based β-hexosaminidase assay (Weatherly et al., 2013), adapted for CPC.

Four antigen doses were assayed: high (0.005 μg/mL), moderate (0.0004 μg/mL), and low (0.0001 and 0.00007 μg/mL). In the absence of CPC, these doses elicited average absolute degranulation responses, respectively, of 51% ± 1 (SEM), 33% ± 2 (SEM), 10.1% ± 0.8% (SEM), and 7.0% ± 0.3% (SEM). Note that, in the absence of IgE sensitization and Ag stimulation, RBL cell spontaneous degranulation is unaffected by CPC doses ranging 10^-6^ to 10 µM (Figure 1E). Thus, Figure 1E suggests that an additional environmental agent, antigen, is needed to elicit degranulation responses.

For IgE-primed RBL mast cells incubated for 1 h in BT containing CPC and the high Ag dose 0.005 µg/mL Ag, CPC significantly inhibits degranulation at 10 µM (Figure 1A). No significant inhibition of degranulation is observed for CPC doses 5 µM and below during the 1 h Ag/CPC co-exposure (Figure 1A). However, when a CPC pretreatment was added, for 30 min prior to the 1 h Ag/CPC co-exposure (P&C), inhibition of degranulation begins at a lower dose, 5 µM CPC (Figure 1A).

When the same co-exposure experiment was conducted instead with a moderate Ag dose, 0.0004 µg/mL Ag, CPC significantly inhibits degranulation in a dose-responsive manner beginning at a much lower dose, 1 µM, (Figure 1B and (Raut et al., 2022)). Upon CPC pretreatment for 30 min prior to the 1 h Ag/CPC exposure (P&C), 5 and 10 µM CPC more strongly inhibit degranulation compared to the samples which had only 1 h Ag/CPC exposure (Figure 1B).

In the presence of low Ag dose stimulation, 0.0001 and 0.00007 µg/mL (Figure 1C &D), CPC also dose-responsively inhibits degranulation, again starting significantly at 1 µM (Figure 1C & D). (Pretreatment experiments, “P&C,” were not conducted with low Ag doses.) By 5 µM, CPC dampens the release of granules down to spontaneous levels (Figure 1C & 1D).

These data show that CPC inhibits RBL mast cell degranulation more potently as the antigen dose decreases. Furthermore, 30 min pre-treatment with CPC, ahead of Ag addition, enhances CPC’s inhibitory effects, which are also clear under conditions 1 h co-exposure to Ag/CPC. We chose to utilize 30 min CPC pre-treatment for each of the measurements of Ca^2+^ levels, which dramatically jump within 5 min of Ag addition, in the three interrogated compartments (ER, Figure 2; mitochondria, Figure 3; cytosol, Figure 4), in order capture any CPC effects that require more than 5 min exposure time.

**Figure 2.**
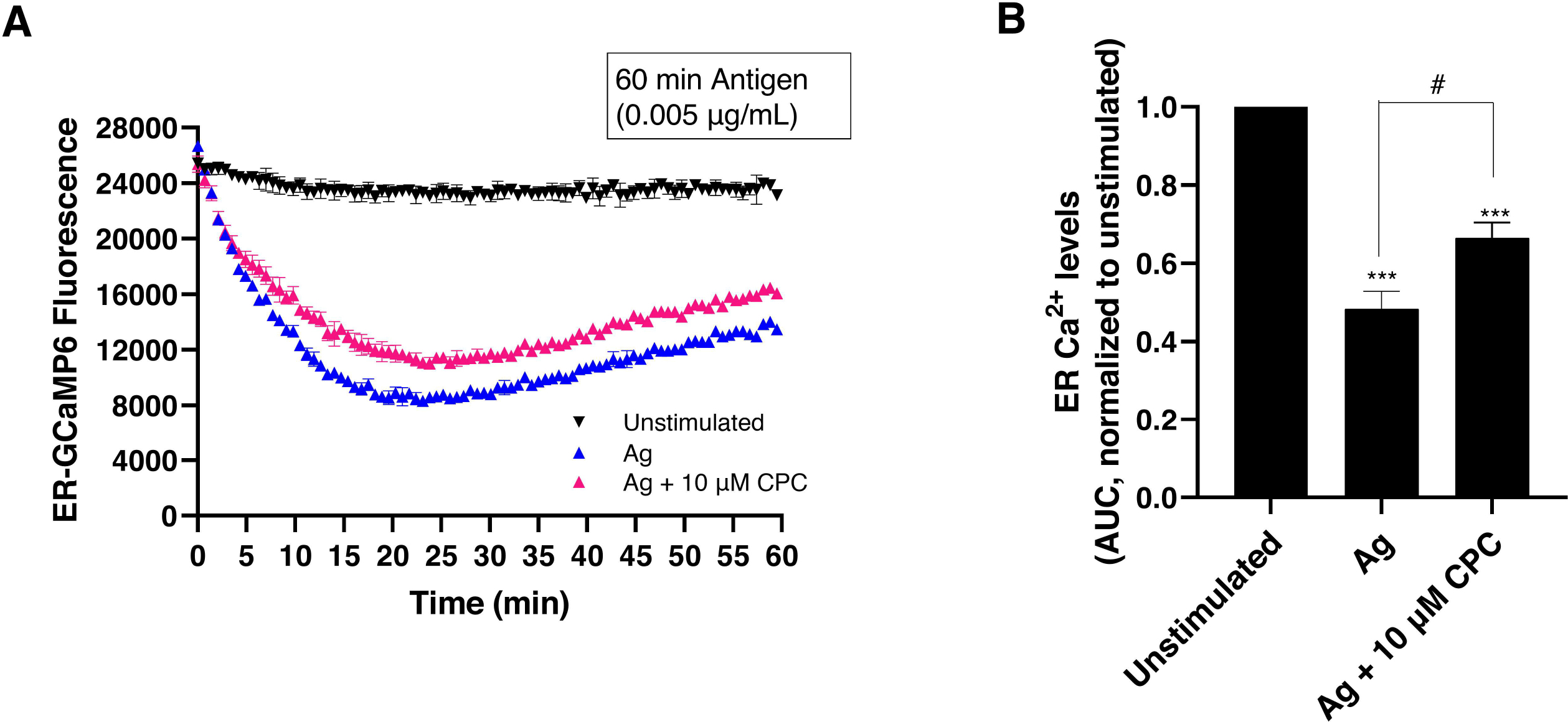
CPC effects on ER Ca^2+^ in Ag-stimulated RBL mast cells. RBL mast cells were transiently transfected with ER-GCaMP6-210 construct, and the ER Ca^2+^ levels were measured using a plate reader. Cells were sensitized with IgE for 1 h, pretreated with 0 or 10 µM CPC Ca^2+^-free BT buffer for 30 min, and exposed to ± 0.005 µg/mL Ag with 0 or 10 µM CPC in Ca^2+^- free BT buffer for 1 h. Representative graphs show ER-GCaMP6-210 fluorescence immediately following Ag/CPC exposure **(A)**. Average background fluorescence from untransfected cells was subtracted from all time points. Area under the curve (AUC) was calculated using the background-subtracted fluorescence time-series graph; these values were then normalized to Unstimulated, 0 µg/mL Ag + 0 µM CPC on that day **(B)**. Values presented in **(A)** are means ± SD for a single experiment with three replicates per treatment. Values presented in **(B)** are means ± SEM of at least three independent experiments, each experiment in triplicate. Statistically significant results are represented by ***p < 0.001 as compared to Unstimulated, and by ^#^p < 0.05 as compared to Ag, as determined by one-way ANOVA followed by Tukey’s post-hoc test.

**Figure 3.**
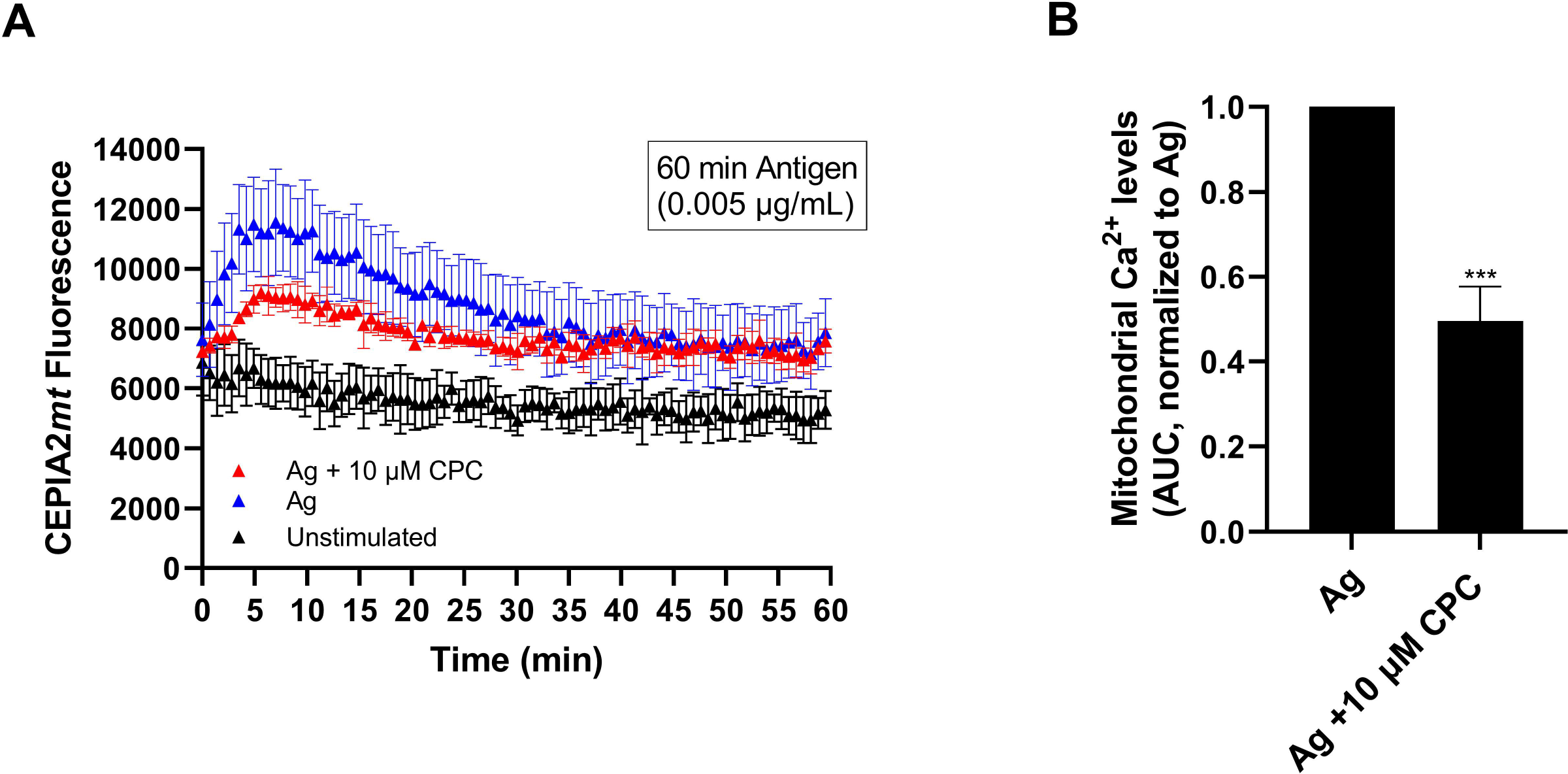
CPC effects on mitochondrial Ca^2+^ in Ag-stimulated RBL mast cells. RBL mast cells were transiently transfected with CEPIA2*mt* construct, and the mitochondrial Ca^2+^ levels were measured using a plate reader. Cells were sensitized with IgE for 1 h, pretreated with 0 or 10 µM CPC in BT buffer for 30 min, and exposed to ± 0.005 µg/mL Ag with 0 or 10 µM CPC in BT buffer for 1 h. Representative graphs show CEPIA2mt fluorescence immediately following Ag/CPC exposure **(A)**; average background fluorescence, from untransfected cells, was subtracted from all time points prior to plotting. Area under the curve (AUC) was calculated using the background-subtracted fluorescence time-series graph; these values were then normalized to Ag + 0 µM CPC on that day **(B)**. Values presented in **(A)** are means ± SD for a single experiment with three replicates per treatment. Values presented in **(B)** are means ± SEM of at least three independent experiments, each experiment was a triplicate. Statistically significant result, as compared to Ag, is represented by ***p < 0.001, determined by an unpaired one-tailed t-test.

**Figure 4.**
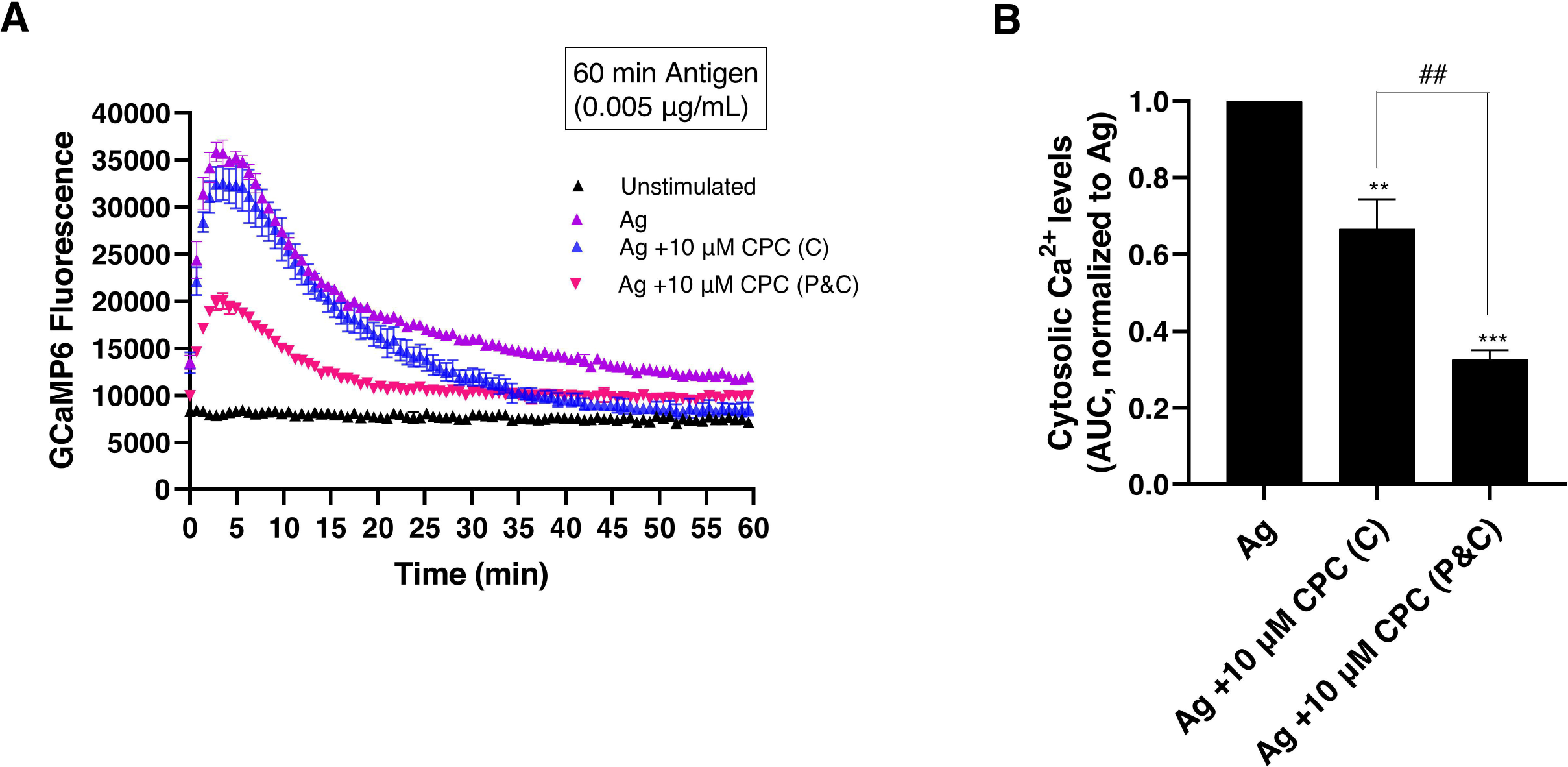
CPC effects on cytosolic Ca^2+^ in Ag-stimulated RBL mast cells. RBL mast cells were transiently transfected with GCaMP6 construct, and the cytosolic Ca^2+^ levels were measured using a plate reader. Cells were sensitized with IgE for 1 h, pretreated with 0 or 10 µM CPC in BT buffer for 30 min, and exposed to ± 0.005 µg/mL Ag and 0 or 10 µM CPC in BT buffer for 1 h. Representative graphs show GCaMP6 fluorescence immediately following Ag/CPC exposure **(A)**; average background fluorescence, from untransfected cells, was subtracted from all time points prior to plotting. Area under the curve (AUC) was calculated using the background-subtracted fluorescence time-series graph; these values were then normalized to Ag + 0 µM CPC on that day **(B)**. Values presented in **(A)** are means ± SD for a single experiment with three replicates per treatment. Values presented in **(B)** are means ± SEM of at least three independent experiments, each experiment was a triplicate. Some samples, denoted “P&C,” in graphs **(A)** and **(B)**, had CPC pretreatment for 30 min prior to the 1 h Ag/CPC exposure; “C” means CPC exposure only during the hour of Ag. Statistically significant results are represented by **p < 0.01 and ***p < 0.001 when compared to Ag-alone and by ^##^ p < 0.01 when compared to Co, determined by one-way ANOVA followed by Tukey’s post-hoc test.

### 3.2. CPC inhibits Ca^2+^ release from the endoplasmic reticulum in RBL mast cells

In Ag-stimulated mast cells, PIP_2_-generated second messenger IP_3_ binds to the IP_3_ receptor on the ER membrane, leading to the depletion of ER Ca^2+^ stores, thus triggering influx of external Ca^2+^ via SOCE (Hogan et al., 2010; Vig et al., 2006). To measure this Ag-stimulated Ca^2+^ release from the ER, RBL cells were transiently transfected via nucleofection with the genetically-encoded Ca^2+^ indicator ER-GCaMP6-210, a fluorescent protein that detects changes in Ca^2+^ levels (de Juan-Sanz et al., 2017). Cells were IgE-sensitized, then pre-treated with 10 μM CPC for 30 min, and then ER-GCaMP6 fluorescence measured in the plate reader immediately following Ag stimulation, in the presence or absence of 10 μM CPC. The AUC is a quantitative measure of the amount of Ca^2+^ in the ER, integrated over the 1 h of measurement. As expected, Ag stimulation triggered an immediate, robust, and continuous decrease in Ca^2+^ levels in the ER for ∼15 minutes (Figure 2A). Refilling of the ER Ca^2+^ stores, indicated by a slow rise in fluorescence during the final ∼half hour, occurs following this decrease (Figure 2A). CPC significantly hampers the efflux of Ca^2+^ from the ER into the cytosol, indicated by the substantially higher AUC of Ag + CPC, 0.67 ± 0.04 (SEM) when compared to Ag alone, 0.48 ± 0.04 (SEM) (Figure 2B).

### 3.3. CPC lowers mitochondrial Ca^2+^ uptake in RBL mast cells

The mitochondria serve as a Ca^2+^ buffer by taking up some of the Ca^2+^ released from the ER (de Brito & Scorrano, 2008). Mitochondrial Ca^2+^ levels were measured via fluorescence of CEPIA2mt, a construct that detects changes in mitochondrial Ca^2+^ levels (Suzuki et al., 2014). Transfected cells were IgE-sensitized, then pre-treated with 10 μM CPC for 30 min, and then CEPIA2mt fluorescence measured in the plate reader immediately following Ag stimulation, in the presence or absence of 10 μM CPC. The AUC is a quantitative measure of the amount of Ca^2+^ in the mitochondria, minus signal from unstimulated cells, integrated over the 1 h of measurement. Following Ag stimulation, there is an immediate rise in CEPIA2mt fluorescence, reaching its peak within the first ∼5 to 10 minutes (Figure 3A). A gradual decrease in Ca^2+^ levels follow the peak, which later reaches a plateau (Figure 3A). CPC significantly decreases the uptake of Ca^2+^ into mitochondria (Figure 3A & 3B). Comparing the AUC value of Ag + CPC to that of Ag alone, CPC decreases mitochondrial Ca^2+^ uptake 0.49-fold ± 0.08 (SEM) (Figure 3B). These data show that CPC strongly inhibits mitochondrial Ca^2+^ uptake in Ag-stimulated RBL cells.

### 3.4. CPC suppresses influx of Ca^2+^ into the cytosol in RBL mast cells

Elevated, sustained levels of cytosolic Ca^2+^ are crucial for degranulation (Holowka et al., 2012). To investigate the effect of CPC on Ag-stimulated Ca^2+^ concentration rise in the cytosol, RBL cells were transiently transfected via nucleofection with the genetically-encoded Ca^2+^ indicator GCaMP6, a fluorescent protein that detects changes in Ca^2+^ levels. There were two groups of CPC treatments, those exposed to 60 mins CPC (“C”) and those exposed to combined 90 mins of CPC (“P&C”). Following pretreatment with ± 10 μM CPC for 30 minutes, GCaMP6 fluorescence was monitored in the plate reader immediately following Ag stimulation ± CPC. The fluorescence data collected was background-subtracted and graphed (Figure 4A), and AUC (Figure 4B) was calculated using the average of the unstimulated group as the y-axis baseline. GCaMP6 fluorescence increased following Ag stimulation (Figure 4A) as expected (Weatherly et al., 2018).

There is a rapid increase in GCaMP6 fluorescence within the first 10 minutes after Ag addition, resulting primarily from ER Ca^2+^ release into the cytosol (Holowka et al., 2012; Mohr & Fewtrell, 1987). CPC suppresses this rapid increase in cytosolic Ca^2+^ (Figure 4A) due to ER depletion (Hogan et al., 2010; Vig et al., 2006), particularly following 30 min CPC pre-treatment (“P&C”). This CPC suppression of the rapid increase in cytosolic Ca^2+^ due to ER Ca^2+^ efflux following Ag stimulation (Figure 4A) aligns with the data showing CPC inhibition of ER Ca^2+^ efflux (Figure 2A & B).

CPC also suppresses the plateau phase (Figure 4A) of sustained Ca^2+^ influx through plasma membrane CRAC channels (Mohr & Fewtrell, 1987). In sum, CPC (10 μM) statistically significantly inhibits the integrated Ag-stimulated cytosolic Ca ^2+^ concentration (Figure 4B). Note that 30 min pre-treatment with CPC (“P&C”) leads to even more significant suppression, decreasing the AUC to 0.32 ± 0.02 (SEM) of the unstimulated control (0 μg/mL Ag + 0 μM CPC) level, as compared to the “C” samples’ dampening of AUC to 0.67 ± 0.08 (SEM) of the unstimulated control (Figure 4B). These data indicate that CPC suppresses calcium mobilization in Ag-stimulated RBL cells. Thus, we next tested whether this effect is partially mediated by CPC inhibition of plasma membrane CRAC channel function by voltage or pH modulation.

### 3.5. CPC does not affect PMP or cytosolic pH in RBL mast cells

To investigate the effect of CPC on plasma membrane potential (PMP) and cytosolic pH, RBL mast cells were transiently transfected with ArcLight-A242, a genetically-encoded voltage indicator that detects changes in both PMP and cytosolic pH (Jin et al., 2012), via nucleofection followed by growth overnight at 37℃ and 5% CO_2_. The use of a likely perturbant-resistant (Brejc et al., 1997; Chalfie & Kain, 2005; Weatherly et al., 2018; Weatherly et al., 2016) fluorescent protein reporter was necessary in this investigation because relevant doses of CPC directly interfere with (dose-responsive increase) the fluorescence of the organic voltage-sensitive dye DiSC_3_(5) (Figure S1). The following day, confocal images were taken for each transfected cell (Figure 5A) at various time points through 30 min during exposure to control (BT) or 10 μM CPC. Average fluorescence around the plasma membrane was quantified using FIJI ImageJ and normalized to the initial time point, 0 min, for each treated cell. If CPC altered cytosolic pH or PMP within 30 min, a change in ArcLight fluorescence would be detected. There was no significant change in ArcLight fluorescence across the various time points following 10 μM CPC exposure compared to the BT control (Figure 5B).

**Figure 5.**
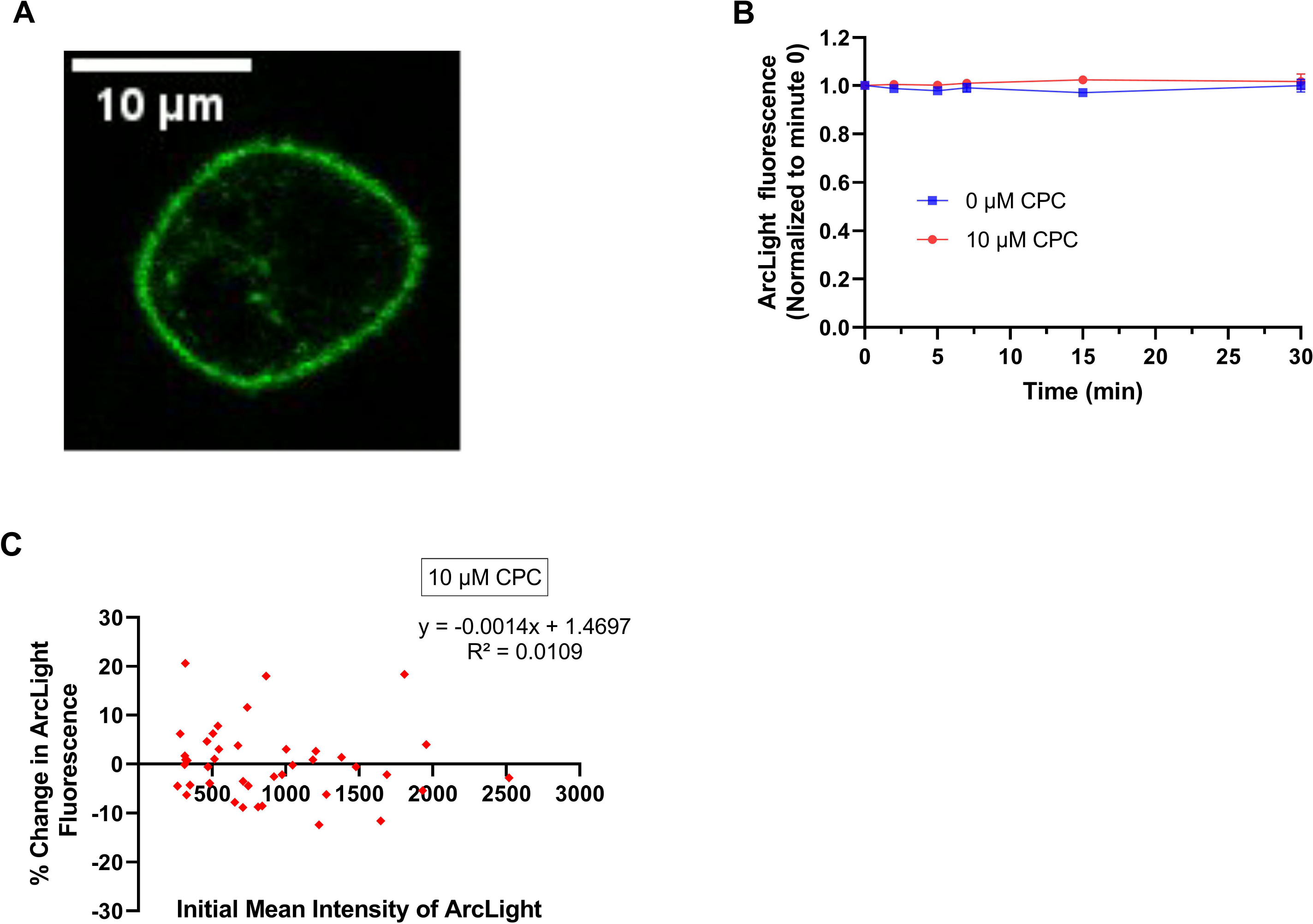
CPC effects on fluorescence of ArcLight-A242 in RBL mast cells. A representative live-cell confocal microscopy image of an RBL mast cell transiently transfected with ArcLight-A242 construct prior to CPC exposure **(A)** (scale bar 10 µm). ArcLight-A242-transfected RBL cells were washed with BT and exposed to control (N=33) or 10 µM CPC (N=41) for 30 min during the live-cell confocal microscopy imaging. The average fluorescence at the plasma membrane at each time point was measured, background-subtracted, and normalized to the 0 min time point of each cell **(B)**. Values presented, in **(B)**, are mean ± SEM of at least 3 independent days of experiment per treatment. The effect of cellular expression level of ArcLight-A242 on CPC effects on its fluorescence is plotted in **(C)**. The percentage change in fluorescence at 30 min time exposure, for each individual cell, is plotted as a function of that cell’s initial mean intensity of ArcLight-A242 for 10 µM CPC-treated cells in **(C)**. The equation of linear regression and R^2^ values are displayed.

To determine if the cellular expression levels (“brightness”) of ArcLight affect its fluorescence change in response to CPC, we plotted each cell’s percentage change in ArcLight fluorescence during the 30 min exposure period against its own initial mean fluorescence intensity of ArcLight (Figure 5C). The linear regression results (Figure 5C) indicate that ArcLight expression level does not correlate with the percentage change in ArcLight fluorescence. Thus, the lack of CPC modulation of ArcLight fluorescence is consistent across a wide range of plasmid expression levels (“brightness”) of ArcLight in cells.

Together, these data indicate that CPC does not interfere Arclight fluorescence and, thus, does not affect PMP or cytosolic pH of RBL cells.

### 3.6. CPC halts microtubule polymerization in living RBL mast cells but not *in vitro*

To test the hypothesis that dampened cytosolic Ca^2+^ mobilization (Figure 4) results in inhibited microtubule polymerization, we transfected RBL mast cells with EGFP-alpha-tubulin, exposed to Ag, and imaged with confocal microscopy (Weatherly et al., 2018). Images of representative cells are shown (Figures 6A-6C). In control, Ag-unstimulated cells, the tubulin construct is localized near the plasma membrane and in a microtubule organization center located near the nucleus (Joshi, 1994) (Figure 6A). Ag stimulation leads to dramatic microtubule filamentation (Figure 6B), while co-exposure with CPC for the 1 h of Ag stimulation negates this Ag effect (Figure 6C). The percentage of cells with microtubule polymerization was determined as detailed in Methods, to quantify these data. We observed a ∼50% decrease in percentage of cells undergoing microtubule polymerization vs. Ag-alone, in cells exposed to 5 μM CPC (Figure 6D), and a total shutdown to Ag-unstimulated levels in cells exposed to 10 μM CPC (Figure 6D).

**Figure 6.**
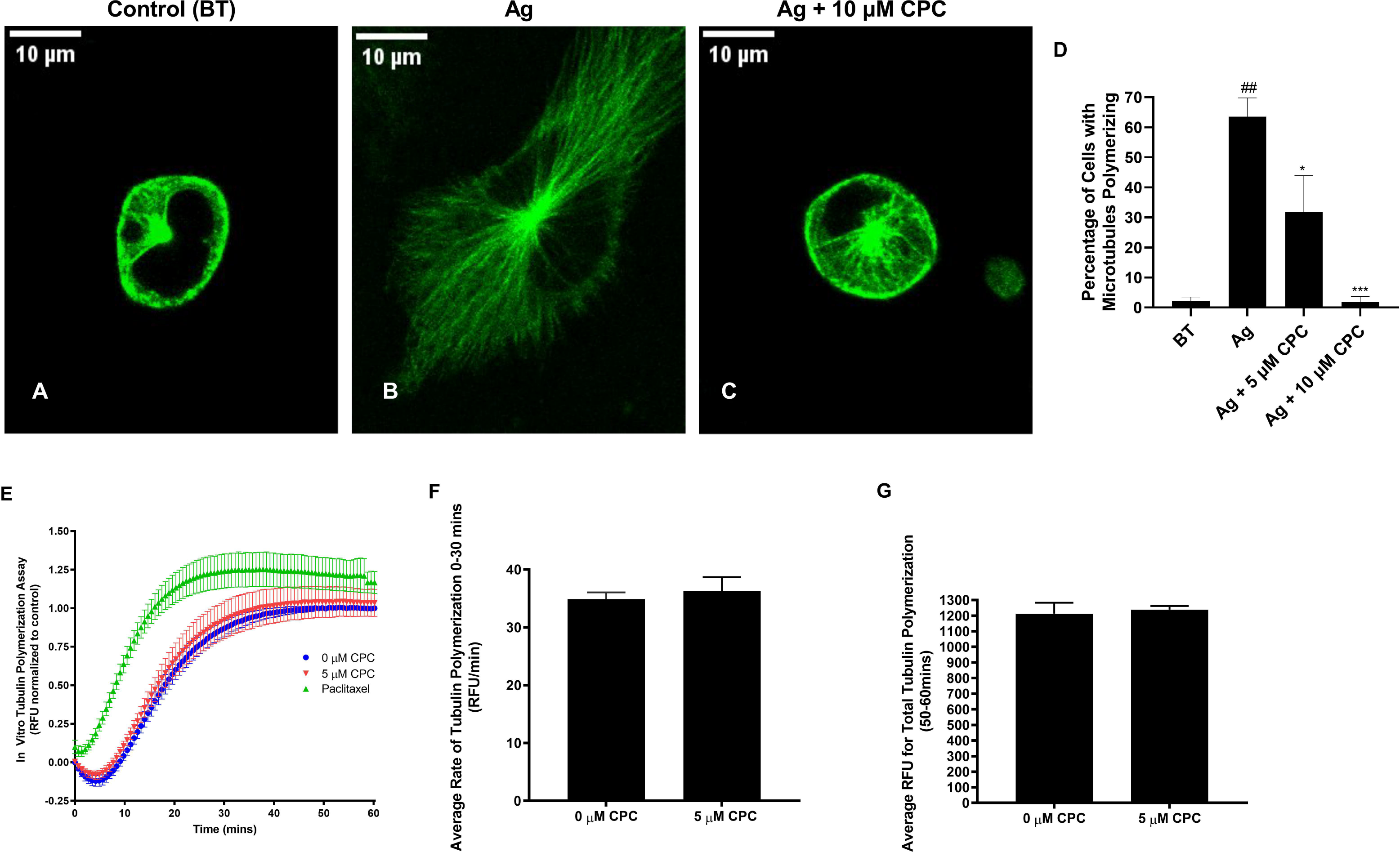
CPC effects on tubulin polymerization in RBL mast cells and *in vitro* assay. RBL mast cells were transiently transfected with EGFP-alpha tubulin construct overnight, washed in BT, and exposed to BT (Control) or 0.005 μg/mL Ag ± 5 µM CPC or 10 µM CPC for 1 h. Confocal microscopy images (scale bar 10 µm) were taken after washing off the CPC with BT. Representative live-cell confocal microscopy images of transfected RBL cells after treatment are shown **(A-C)**. The percentage of cells with microtubule polymerization was calculated by manual classification as in Methods **(D)**. Values presented are mean ± SEM of at least three independent experiments and are derived from analysis of at least 38 cells per treatment. Statistically significant results, as compared to control (BT) are represented by ^##^p < 0.001, and as compared to Ag by *p < 0.05 and **p < 0.001, determined by one-way ANOVA followed by Tukey’s post-hoc test. Using an *in vitro* assay, the direct effect of CPC on tubulin polymerization was measured **(E-G)**. Duplicated-averaged, background-subtracted fluorescence intensity for tubulin polymerization was normalized to the final time point of the 0 μM CPC control (y-axis) and plotted for 0 μM CPC group, 5 μM CPC group, and a paclitaxel positive-control group against time for 1 hour **(E)**. Values presented in **(E)** are means ± SEM of 4 independent experiments. The rate of tubulin polymerization (y-axis) of 0 μM CPC and 5 μM CPC was measured over the first 30 min of tubulin polymerization **(F)**. Data in **(F)** represent mean slope of fluorescence intensity ± SD of 4 independent experiments performed in duplicate, probed by two-tailed t-test (no significance). Total tubulin polymerization was assessed in **(G)**. Average fluorescence intensity (y-axis, **G**) was measured in the final 10 min of tubulin polymerization for 0 μM CPC and 5 μM CPC groups. Data represent mean fluorescence intensity ± SD of 4 independent experiments performed in duplicate, probed by two-tailed t-test (no significance).

An *in vitro* assay was performed to test the hypothesis that CPC directly binds to and inhibits tubulin polymerization (Figures 6E-6G). We first performed a control experiment to test whether CPC (5 μM) interferes with the fluorescence of the proprietary dye employed within this kit; CPC does not interfere (Figure S2). Fluorescence intensity of tubulin polymerization over one hour suggests no CPC effects (Figure 6E) whereas clear positive-stimulatory effects of the drug paclitaxel are measured and plotted for comparison (Figure 6E). To quantitate these results, the initial (first 30 min) rates of tubulin polymerization were calculated and were found to be not significantly different between the 0 μM control group and the 5 μM CPC group (Figure 6F). Furthermore, the total amounts of tubulin polymerization, assessed over the final 10 minutes, between the 0 μM control group and the 5 μM CPC group, were not significantly different (Figure 6G). These *in vitro* tubulin findings indicate that CPC does not abolish tubulin polymerization via a direct mechanism of binding to tubulin protein or to its nucleotide activator GTP.

## 4. Discussion

Despite widespread, high-concentration usage for decades, science and society have largely ignored CPC’s effects on any eukaryote. While an antimicrobial agent charged with protecting human health from microbes, CPC also potently inhibits immune cell signaling (Raut et al., 2022), at doses ∼1000-fold lower than those used in personal care and food products. Notably, CPC is more potent at inhibiting MC function than the banned/removed (Gottlieb, 2019; Kux, 2017) antimicrobial agent triclosan, with 5 µM triclosan causing ∼20% inhibition of degranulation (Weatherly et al., 2013) compared to ∼50% inhibition caused by 5 µM CPC (Figure 1B) with the same Ag stimulation and experimental conditions.

In this study, we discovered a CPC mode of action, inhibition of Ca^2+^ mobilization. Thus, CPC may alter Ca^2+^-dependent signal transduction pathways in numerous eukaryotic cell types including T cells (Marano et al., 1993; Trebak & Kinet, 2019).

CPC inhibits the degranulation of mast cells stimulated by a low, moderate, and high Ag concentrations in a dose- and timing-responsive manner at non-cytotoxic doses. CPC is a more potent inhibitor at longer exposure times (extra 30 min; Figure 1A & B) and at lower Ag concentrations (Figure 1C & D).

CPC inhibits mitochondrial function (Chávez & Bravo, 1982; Datta et al., 2017), estrogen signaling (Datta et al., 2017) and dopamine D3 receptors (D3R) (Pottel et al., 2020); CPC also produces oxidative stress (Park et al., 2016). These biochemical effects could be potentially involved in the observed MC inhibition of CPC. However, CPC likely does not hamper MC function (Figure 1) via mitochondrial dysfunction because these degranulation assays are carried out in glucose, in the presence of which CPC does not inhibit mitochondrial ATP production (Weller et al., 2022). Estrogens are known to enhance MC degranulation (Narita et al., 2007). However, CPC likely does not inhibit MC function via estrogen disruption because the relatively low stimulation of MC degranulation by estrogens and reversal thereof by estrogen blockers (Narita et al., 2007) is a relatively a low effect compared to the potency of Ag/CPC (Figure 1). CPC induced reactive oxygen species (ROS) production in amphibian embryos (Park et al., 2016). However, ROS production is actually required for degranulation (Swindle et al., 2004), so CPC stimulation of ROS is likely not the mechanism underlying CPC MC inhibition. Also, CPC inhibition of degranulation is likely not due to its inhibition of D3R because lowered D3R function correlates with increased MC function (Xue et al., 2018). Thus, a separate molecular mechanism likely underlies CPC effects on MC.

Our process of deciphering CPC’s mode of action began with the observation that CPC is a disruptor of interactions between PIP_2_ and several PIP_2_-binding proteins (Raut et al., 2022). Because PIP_2_ is a critical player in SOCE, we then hypothesized that CPC disruption of the enzymatic cleavage of PIP_2_ could cause SOCE inhibition and, thus, degranulation inhibition. Therefore, we hypothesized that CPC interaction with PIP_2_ or with PIP_2_-binding partners would disrupt the production of IP_3_ and thereby lead to decreased Ca^2+^ efflux from the ER. Thus, we first tested CPC effects on levels of Ca^2+^ in the ER lumen. Indeed, CPC inhibits the efflux of Ca^2+^ from the ER lumen into the cytosol (Figure 2). Future planned experiments include use of an IP_3_ sensor (Matsu-Ura et al., 2019) to directly measure CPC effects on the levels of this second messenger.

While there exist data that PIP_2_ directly complexes with both ER and plasma membrane Ca^2+^ channels, these data show an inhibitory effect of PIP_2_ on these channels’ functioning (Kaznacheyeva et al., 2000; Lupu et al., 1998). Since CPC is a known direct disruptor of PIP_2_ (Raut et al., 2022), we postulate that the IP_3_ disruption hypothesis is a more likely mechanism than a direct CPC interference in PIP_2_-Ca^2+^ channel interactions (which would actually be stimulatory).

In addition to being responsible for IP_3_ production, PIP_2_ also assists with soluble N-ethylmaleimide-sensitive factor attachment protein receptor (SNARE) machinery needed for granule fusion to the PM (Dai et al., 2007; Paddock et al., 2008; Tadokoro et al., 2015) and other numerous crucial roles in the cell via binding to and regulation of cytoskeletal elements (Janmey et al., 2018). So even though our immediate hypothesis is that CPC disrupts the production of IP_3_ from PIP_2_, as an explanation for all the biochemical sequelae of that effect, there are numerous other possible biochemical effects on mast cell signaling that could be caused by CPC disruption of PIP_2_. This area will be subject to future investigation.

Flow of Ca^2+^ through the ER, mitochondria, and cytosol is orchestrated in the cell, so CPC inhibition of one compartment’s Ca^2+^ levels affects the others. Depletion of Ca^2+^ in the ER lumen activates the CRAC channel via STIM1 (Putney, 1986). CPC inhibition of Ca^2+^ efflux from the ER (Figure 2), suggests STIM1 should be activated to a lesser degree in the presence of CPC, leading to reduced Ag-stimulated cytosolic Ca^2+^ levels, as we found in Figure 4. Also, less Ca^2+^ leaving the ER or inflowing through CRAC channels should result in less Ca^2+^ taken up into the mitochondria, as recorded in Figure 3.

While CPC disruption of PIP_2_-dependent IP_3_ production could explain the Ca^2+^ mobilization inhibition (Figures. 2, 3, 4), there are other potential explanatory mechanisms. For example, chemical depolarization of the plasma membrane, detected as a change in PMP, can cause SOCE inhibition in mast cells (Mohr & Fewtrell, 1987). Thus, using ArcLight A-242, we tested whether CPC affects PMP: CPC does not interfere with the integrity of the PMP (Figure 5). ArcLight A-242, with a pHlourin domain, simultaneously detects cytosolic pH near the PM. Acidification of the cytoplasm inhibits CRAC channels (Beck et al., 2014; Sangroula et al., 2020) and blocks the binding of IP_3_ to its ER receptor, thus inhibiting SOCE (Tsukioka et al., 1994). Our results indicate that CPC does not cause acidification of the cytosol (Figure 5). Thus, CPC does not inhibit mast cell function via either PMP disruption or acidification of the cytoplasm. Thus, we instead hypothesize that CPC may act to inhibit Ca^2+^ mobilization via interference with Ag-stimulated phosphorylation events, including PLCγ which hydrolyzes PIP_2_ into IP_3_ (Kinet, 1999), necessary for SOCE. These questions will be addressed in future experiments.

Polymerized microtubules are needed to transport granules to the PM for degranulation (Guo et al., 1998; Smith et al., 2003). Microtubule polymerization requires high cytosolic Ca^2+^ levels, which stimulate the positive tubulin regulator Git1 (Sulimenko et al., 2015). In this study, we have shown that CPC inhibits Ag-stimulated microtubule polymerization in cells (Figure 6). This effect could be a direct consequence of CPC’s suppression of Ag-stimulated cytosolic Ca^2+^ levels (Figure 4). Our results are buttressed by a recent high-throughput comparative study (Pottel et al., 2020). While this study did not directly measure CPC effects on microtubules (such as tubulin gene or protein expression or microtubule polymerization / cellular distribution), the authors employed the BioMAP Diversity PLUS phenotypic profiling system to generate data and computationally compare it to libraries of compounds with known modes of action (Kleinstreuer et al., 2014). BioMAP Diversity PLUS uses multiple systems consisting of primary human cells stimulated by various cytokines and measures endpoints like cytotoxicity, proliferation, and protein (cell adhesion molecules, cytokines, etc) levels via ELISA (Kleinstreuer et al., 2014). In this indirect though suggestive study, CPC was found to be phenotypically similar, in the combination of endpoints assessed, to a known microtubule disruptor (Pottel et al., 2020).

While CPC inhibits microtubule polymerization in cells (Figure 6), a separate *in vitro* tubulin assay revealed that CPC actually does not inhibit microtubule polymerization by direct binding/interference with tubulin or with the nucleotide guanosine-5’-triphosphate (GTP) required for microtubule assembly (Arai & Kaziro, 1977) (Figure 6 E-G). The lack of a CPC effect on the *in vitro* rate of tubulin polymerization (Figure 6F) or total tubulin polymerization (Figure 6G) further implicates CPC’s inhibition of cytosolic Ca^2+^ influx as the biochemical mechanism underlying CPC’s microtubule disruption.

CPC inhibition of mast cell signaling may extend to other cell types, such as T cells, that utilize similar signaling leading to SOCE: T-cell receptor (TCR) activation (Kuby, 1997; Smith-Garvin et al., 2009) also leads to PLCγ activation, IP_3_ generation, and SOCE (Dolmetsch & Lewis, 1994; Roos et al., 2005), as in mast cells. Because signaling upstream of SOCE is very similar between mast cells and T-cells, our findings of CPC inhibition of mast cell signaling may be predictive of T cell toxicity from this drug.

In conclusion, we report CPC’s inhibition of mast cell function and the underlying biochemical mechanisms. We have shown that CPC inhibits degranulation at doses ∼1000-fold lower than those used in consumer products. CPC inhibits mast cell signaling/degranulation (Figure 7) by inhibiting SOCE via suppression of Ca^2+^ efflux from the ER and of Ca^2+^ uptake into both the cytosol and the mitochondria. This SOCE disruption is not due to CPC interference with PMP or cytosolic pH. Additionally, CPC inhibits Ca^2+^ dependent microtubule polymerization in cells, but not via direct tubulin binding/interference. In summary, we show that CPC is a signaling toxicant that is more potent than the banned/removed drug triclosan and that its mode of action involves a core signaling process, SOCE, used by numerous physiological processes. Any cell type that utilizes Ca^2+^ signaling may be susceptible to CPC toxicity (Feske, 2007; Hill-Eubanks et al., 2011).

**Figure 7.**
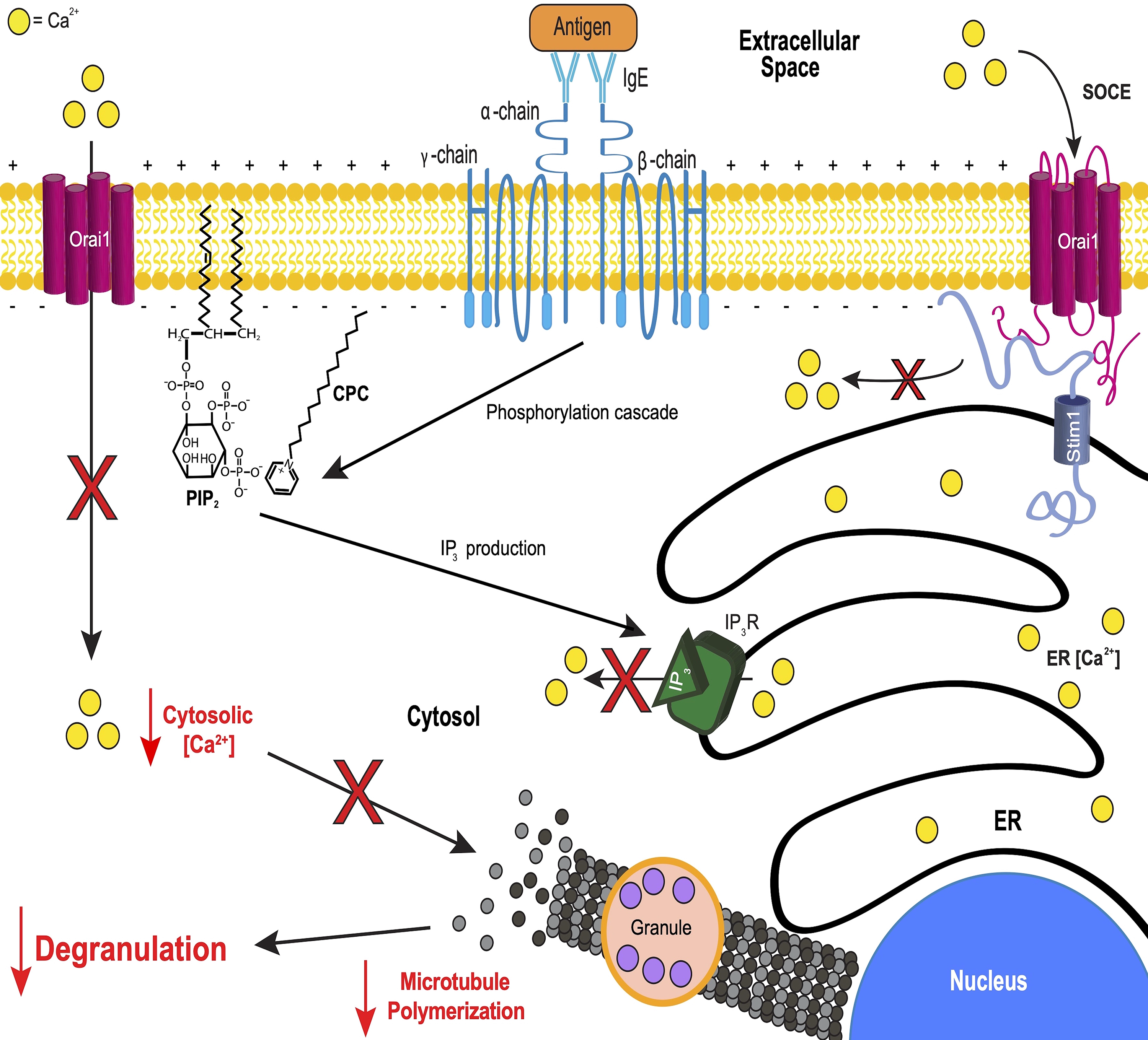
Summary of CPC effects on mast cell signaling. Antigen cross-linking of IgE-bound FcεRI receptors leads to early phosphorylation events, causing the enzymatic cleavage of PIP_2_ to create IP_3_, which binds to its receptor on the ER membrane, initiating dynamic mobilization of calcium from the ER into the cytosol. This ER Ca^2+^ mobilization is hampered in CPC-treated cells. In healthy cells, depletion of ER Ca^2+^ stores stimulate Stim1-Orai1 interaction and subsequent activation of SOCE, leading to mass influx of extracellular Ca^2+^ into the cytosol. However, CPC treatment diminishes this Ca^2+^ influx, albeit not via CPC effects on PMP or on cytosolic pH. Mass Ca^2+^ influx is needed to trigger microtubule polymerization, leading to mast cell degranulation, but CPC abolishes both microtubule polymerization in the cell and inhibits subsequent degranulation. This CPC effect on microtubules is indirect, not a direct binding of CPC to tubulin. Interference by CPC on key signaling events leading to inhibition of degranulation response is illustrated by red X or arrow symbols. (The size of the lipids in this figure have been increased for emphasis.)

## Abbreviation

Ag: antigen
ATP: adenosine triphosphate
BSA: bovine serum albumin
BT: Tyrodes-bovine serum albumin
CMC: critical micelle concentration
CPC: cetylpyridinium chloride
CRAC: Ca^2+^ release-activated Ca^2+^
D3R: Dopamine D3 receptor
DNP: anti-dinitrophenyl
ER: endoplasmic reticulum
GEVI: genetically-encoded voltage indicators
GTP: guanosine-5’-triphosphate
IgE: immunoglobulin E
IP_3_: inositol 1,4,5-triphosphate
LDH: lactate dehydrogenase
MARCKS: myristoylated alanine-rich C-kinase substrate
MC: mast cell
MCU: mitochondrial Ca^2+^ uniporter
PIP_2_: phosphatidylinositol 4,5-bisphosphate
PLCγ: phospholipase C gamma
PM: plasma membrane
PMP: plasma membrane potential
RBL-2H3: rat basophilic leukemia cells, clone 2H3
ROS: reactive oxygen species
SEM: standard error of the mean
SARS-CoV-2: severe acute respiratory syndrome coronavirus 2
SOCE: store-operated Ca^2+^ entry
STIM1: stromal interaction molecule 1.

## Supporting information

This article contains supporting information.

## Acknowledgments

We thank Dr. Robert Wheeler and members of the Wheeler lab, particularly Siham Hattab and Bailey Blair, for the use of and help with the ibidi heating system and confocal. We are grateful to Dr. Joshua Kelley for help with image analysis and to Marissa Paine for help with lab management. We are also grateful to Dylan Wagner for editing the manuscript.

## References

Abezgauz, L., Kuperkar, K., Hassan, P. A., Ramon, O., Bahadur, P., & Danino, D. (2010). Effect of Hofmeister anions on micellization and micellar growth of the surfactant cetylpyridinium chloride. J Colloid Interface Sci, 342(1), 83–92. https://doi.org/10.1016/j.jcis.2009.08.045

Abramson, J., & Pecht, I. (2007). Regulation of the mast cell response to the type 1 Fc epsilon receptor. Immunol Rev, 217, 231–254. https://doi.org/10.1111/j.1600-065X.2007.00518.x

Alemany, A., Perez-Zsolt, D., Raïch-Regué, D., Muñoz-Basagoiti, J., Ouchi, D., Laporte-Villar, C., Baro, B., Henríquez, N., Prat, N., Gianinetto, M. O., Gutiérrez, M. V., Sánchez-Paniagua, M. G., Henríquez, N. L., Vicente, J. M., Ara, J., Rodriguez-Arias, M. A., Puig, J., Blanco, I., Lopez, C. C., . . . Mitjà, O. (2022). Cetylpyridinium Chloride Mouthwash to Reduce Shedding of Infectious SARS-CoV-2: A Double-Blind Randomized Clinical Trial. Journal of Dental Research, 101(12), 1450–1456. https://doi.org/10.1177/00220345221102310

Alsaleh, N. B., Persaud, I., & Brown, J. M. (2016). Silver Nanoparticle-Directed Mast Cell Degranulation Is Mediated through Calcium and PI3K Signaling Independent of the High Affinity IgE Receptor. PLoS One, 11(12), e0167366. https://doi.org/10.1371/journal.pone.0167366

American Chemical Society. (2021). Molecule of the Week Archive Cetylpyridinium chloride. https://www.acs.org/content/acs/en.html

Arai, T., & Kaziro, Y. (1977). Role of GTP in the assembly of microtubules. J Biochem, 82(4), 1063–1071. https://doi.org/10.1093/oxfordjournals.jbchem.a131777

Beck, A., Fleig, A., Penner, R., & Peinelt, C. (2014). Regulation of endogenous and heterologous Ca²⁺ release-activated Ca²⁺ currents by pH. Cell Calcium, 56(3), 235–243. https://doi.org/10.1016/j.ceca.2014.07.011

Bernauer, U., C.P., Degen G, Dusinska M, Lilienblum W, Nielsen E, Rastogi SC, Rousselle C, & van Benthem J. (2015). Opinion on cetylpyridinium chloride - submission II SCCS (Scientific Committee on Consumer In E. Union (Ed.), (pp. 97).

Berridge, M. J. (1993). Inositol trisphosphate and calcium signalling. Nature, 361(6410), 315–325. https://doi.org/10.1038/361315a0

Blank, U., & Benhamou, M. (2013). Deciphering new molecular mechanisms of mast cell activation. Front Immunol, 4, 100. https://doi.org/10.3389/fimmu.2013.00100

Bonesvoll, P., & Gjermo, P. (1978). A comparision between chlorhexidine and some quaternary ammonium compounds with regard to retention, salivary concentration and plaque-inhibiting effect in the human mouth after mouth rinses. Arch Oral Biol, 23(4), 289–294. https://doi.org/10.1016/0003-9969(78)90021-3

Brejc, K., Sixma, T. K., Kitts, P. A., Kain, S. R., Tsien, R. Y., Ormö, M., & Remington, S. J. (1997). Structural basis for dual excitation and photoisomerization of the Aequorea victoria green fluorescent protein. Proc Natl Acad Sci U S A, 94(6), 2306–2311. https://doi.org/10.1073/pnas.94.6.2306

Chalfie, & Kain. (2005). Green Fluorescent Protein:L Properties, Applications, and Protocols (2 ed.). Wiley.

Chávez, E., & Bravo, C. (1982). Anisotropic action of cetyl pyridinium chloride on rat heart mitochondria. Arch Biochem Biophys, 213(1), 81–86. https://doi.org/10.1016/0003-9861(82)90442-8

Chen, T. W., Wardill, T. J., Sun, Y., Pulver, S. R., Renninger, S. L., Baohan, A., Schreiter, E. R., Kerr, R. A., Orger, M. B., Jayaraman, V., Looger, L. L., Svoboda, K., & Kim, D. S. (2013). Ultrasensitive fluorescent proteins for imaging neuronal activity. Nature, 499(7458), 295–300. https://doi.org/10.1038/nature12354

Clapham, D. E. (1995). Intracellular calcium. Replenishing the stores. Nature, 375(6533), 634–635. https://doi.org/10.1038/375634a0

Dai, H., Shen, N., Araç, D., & Rizo, J. (2007). A quaternary SNARE-synaptotagmin-Ca2+-phospholipid complex in neurotransmitter release. J Mol Biol, 367(3), 848–863. https://doi.org/10.1016/j.jmb.2007.01.040

Datta, S., He, G., Tomilov, A., Sahdeo, S., Denison, M. S., & Cortopassi, G. (2017). In Vitro Evaluation of Mitochondrial Function and Estrogen Signaling in Cell Lines Exposed to the Antiseptic Cetylpyridinium Chloride. Environ Health Perspect, 125(8), 087015. https://doi.org/10.1289/ehp1404

de Brito, O. M., & Scorrano, L. (2008). Mitofusin 2 tethers endoplasmic reticulum to mitochondria. Nature, 456(7222), 605–610. https://doi.org/10.1038/nature07534

de Juan-Sanz, J., Holt, G. T., Schreiter, E. R., de Juan, F., Kim, D. S., & Ryan, T. A. (2017). Axonal Endoplasmic Reticulum Ca(2+) Content Controls Release Probability in CNS Nerve Terminals. Neuron, 93(4), 867–881.e866. https://doi.org/10.1016/j.neuron.2017.01.010

Dolmetsch, R. E., & Lewis, R. S. (1994). Signaling between intracellular Ca2+ stores and depletion-activated Ca2+ channels generates [Ca2+]i oscillations in T lymphocytes. J Gen Physiol, 103(3), 365–388. https://doi.org/10.1085/jgp.103.3.365

Dong, K., Li, L., Chen, C., Tengbe, M. S., Chen, K., Shi, Y., Wu, X., & Qiu, X. (2022). Impacts of cetylpyridinium chloride on the behavior and brain neurotransmitter levels of juvenile and adult zebrafish (Danio rerio). Comp Biochem Physiol C Toxicol Pharmacol, 259, 109393. https://doi.org/10.1016/j.cbpc.2022.109393

Feske, S. (2007). Calcium signalling in lymphocyte activation and disease. Nat Rev Immunol, 7(9), 690–702. https://doi.org/10.1038/nri2152

Food and Drug Administration. (2023). Secondary direct food additives permitted in food for human consumption. Retrieved from https://www.accessdata.fda.gov/scripts/cdrh/cfdocs/cfcfr/CFRSearch.cfm?fr=173.375

Furuno, T., Shinkai, N., Inoh, Y., & Nakanishi, M. (2015). Impaired expression of the mitochondrial calcium uniporter suppresses mast cell degranulation. Mol Cell Biochem, 410(1-2), 215–221. https://doi.org/10.1007/s11010-015-2554-4

Gibson, T. J., Hyvönen, M., Musacchio, A., Saraste, M., & Birney, E. (1994). PH domain: the first anniversary. Trends Biochem Sci, 19(9), 349–353. https://doi.org/10.1016/0968-0004(94)90108-2

Gottlieb, S. (2019). Safety and Effectiveness of Consumer Antiseptic Rubs; Topical Antimicrobial Drug Products for Over-the-Counter Human Use. Federal Register, 84(71). https://www.govinfo.gov/content/pkg/FR-2019-04-12/pdf/2019-06791.pdf

Guo, Z., Turner, C., & Castle, D. (1998). Relocation of the t-SNARE SNAP-23 from lamellipodia-like cell surface projections regulates compound exocytosis in mast cells. Cell, 94(4), 537–548. https://doi.org/10.1016/s0092-8674(00)81594-9

Hill-Eubanks, D. C., Werner, M. E., Heppner, T. J., & Nelson, M. T. (2011). Calcium signaling in smooth muscle. Cold Spring Harb Perspect Biol, 3(9), a004549. https://doi.org/10.1101/cshperspect.a004549

Hogan, P. G., Lewis, R. S., & Rao, A. (2010). Molecular basis of calcium signaling in lymphocytes: STIM and ORAI. Annu Rev Immunol, 28, 491–533. https://doi.org/10.1146/annurev.immunol.021908.132550

Holowka, D., Calloway, N., Cohen, R., Gadi, D., Lee, J., Smith, N. L., & Baird, B. (2012). Roles for ca(2+) mobilization and its regulation in mast cell functions. Front Immunol, 3, 104. https://doi.org/10.3389/fimmu.2012.00104

Hrubec, T. C., Seguin, R. P., Xu, L., Cortopassi, G. A., Datta, S., Hanlon, A. L., Lozano, A. J., McDonald, V. A., Healy, C. A., Anderson, T. C., Musse, N. A., & Williams, R. T. (2021). Altered toxicological endpoints in humans from common quaternary ammonium compound disinfectant exposure. Toxicol Rep, 8, 646–656. https://doi.org/10.1016/j.toxrep.2021.03.006

Hutchinson, L. M., Trinh, B. M., Palmer, R. K., Preziosi, C. A., Pelletier, J. H., Nelson, H. M., & Gosse, J. A. (2011). Inorganic arsenite inhibits IgE receptor-mediated degranulation of mast cells. J Appl Toxicol, 31(3), 231–241. https://doi.org/10.1002/jat.1585

Janmey, P. A., Bucki, R., & Radhakrishnan, R. (2018). Regulation of actin assembly by PI(4,5)P2 and other inositol phospholipids: An update on possible mechanisms. Biochem Biophys Res Commun, 506(2), 307–314. https://doi.org/10.1016/j.bbrc.2018.07.155

Jin, L., Han, Z., Platisa, J., Wooltorton, J. R., Cohen, L. B., & Pieribone, V. A. (2012). Single action potentials and subthreshold electrical events imaged in neurons with a fluorescent protein voltage probe. Neuron, 75(5), 779–785. https://doi.org/10.1016/j.neuron.2012.06.040

Joshi, H. C. (1994). Microtubule organizing centers and gamma-tubulin. Curr Opin Cell Biol, 6(1), 54–62. https://doi.org/10.1016/0955-0674(94)90116-3

Kaznacheyeva, E., Zubov, A., Nikolaev, A., Alexeenko, V., Bezprozvanny, I., & Mozhayeva, G. N. (2000). Plasma membrane calcium channels in human carcinoma A431 cells are functionally coupled to inositol 1,4,5-trisphosphate receptor-phosphatidylinositol 4,5-bisphosphate complexes. J Biol Chem, 275(7), 4561-4564. https://doi.org/10.1074/jbc.275.7.4561

Kennedy, R. H., Pelletier, J. H., Tupper, E. J., Hutchinson, L. M., & Gosse, J. A. (2012). Estrogen mimetic 4-tert-octylphenol enhances IgE-mediated degranulation of RBL-2H3 mast cells. J Toxicol Environ Health A, 75(24), 1451–1455. https://doi.org/10.1080/15287394.2012.722184

Keown, M. B., Henry, A. J., Ghirlando, R., Sutton, B. J., & Gould, H. J. (1998). Thermodynamics of the interaction of human immunoglobulin E with its high-affinity receptor Fc epsilon RI. Biochemistry, 37(25), 8863–8869. https://doi.org/10.1021/bi972354h

Kim, H., Yoo, J., Lim, Y. M., Kim, E. J., Yoon, B. I., Kim, P., Yu, S. D., Eom, I. C., & Shim, I. (2021). Comprehensive pulmonary toxicity assessment of cetylpyridinium chloride using A549 cells and Sprague-Dawley rats. J Appl Toxicol, 41(3), 470–482. https://doi.org/10.1002/jat.4058

Kim, Y. J., Rossa, C., Jr., & Kirkwood, K. L. (2005). Prostaglandin production by human gingival fibroblasts inhibited by triclosan in the presence of cetylpyridinium chloride. J Periodontol, 76(10), 1735–1742. https://doi.org/10.1902/jop.2005.76.10.1735

Kinet, J. P. (1999). The high-affinity IgE receptor (Fc epsilon RI): from physiology to pathology. Annu Rev Immunol, 17, 931–972. https://doi.org/10.1146/annurev.immunol.17.1.931

Kleinstreuer, N. C., Yang, J., Berg, E. L., Knudsen, T. B., Richard, A. M., Martin, M. T., Reif, D. M., Judson, R. S., Polokoff, M., Dix, D. J., Kavlock, R. J., & Houck, K. A. (2014). Phenotypic screening of the ToxCast chemical library to classify toxic and therapeutic mechanisms. Nat Biotechnol, 32(6), 583–591. https://doi.org/10.1038/nbt.2914

Kozak, J. A., and Putney J. W., Jr. (2018). In J. A. Kozak & J. W. Putney, Jr. (Eds.), Calcium Entry Channels in Non-Excitable Cells. CRC Press/Taylor & Francis © 2017 by Taylor & Francis Group, LLC. https://doi.org/10.1201/9781315152592

Krystel-Whittemore, M., Dileepan, K. N., & Wood, J. G. (2015). Mast Cell: A Multi-Functional Master Cell. Front Immunol, 6, 620. https://doi.org/10.3389/fimmu.2015.00620

Kuby, J. (1997). Immunology (Sixth ed.). W.H Freeman, New York.

Kux, L. (2017). Safety and Effectiveness of Health Care Antiseptics; Topical Antimicrobial Drug Products for Over-the-Counter Human Use Federal Register, 82(243). https://www.govinfo.gov/content/pkg/FR-2017-12-20/pdf/2017-27317.pdf

Lin, G. H., Voss, K. A., & Davidson, T. J. (1991). Acute inhalation toxicity of cetylpyridinium chloride. Food Chem Toxicol, 29(12), 851–854. https://doi.org/10.1016/0278-6915(91)90113-l

Lupu, V. D., Kaznacheyeva, E., Krishna, U. M., Falck, J. R., & Bezprozvanny, I. (1998). Functional coupling of phosphatidylinositol 4,5-bisphosphate to inositol 1,4,5-trisphosphate receptor. J Biol Chem, 273(23), 14067-14070. https://doi.org/10.1074/jbc.273.23.14067

Mandal, A. B., Nair, B.U.,. (1991). Cyclic voltammetric technique for the determination of the critical micelle concentration of surfactants, self-diffusion coefficient of micelles, and partition coefficient of an electrochemical probe. The Journal of Physical Chemistry,(95), 9008-9013. https://doi.org/doi.org/10.1021/j100175a106

Mao, X., Auer, D. L., Buchalla, W., Hiller, K. A., Maisch, T., Hellwig, E., Al-Ahmad, A., & Cieplik, F. (2020). Cetylpyridinium Chloride: Mechanism of Action, Antimicrobial Efficacy in Biofilms, and Potential Risks of Resistance. Antimicrob Agents Chemother, 64(8), e00576–00520. https://doi.org/10.1128/AAC.00576-20

Marano, N., Liotta, M. A., Slattery, J. P., Holowka, D., & Baird, B. (1993). Fc epsilon RI and the T cell receptor for antigen activate similar signalling pathways in T cell-RBL cell hybrids. Cell Signal, 5(2), 155–167. https://doi.org/10.1016/0898-6568(93)90067-v

Martin-Verdeaux, S., Pombo, I., Iannascoli, B., Roa, M., Varin-Blank, N., Rivera, J., & Blank, U. (2003). Evidence of a role for Munc18-2 and microtubules in mast cell granule exocytosis. J Cell Sci, 116(Pt 2), 325–334. https://doi.org/10.1242/jcs.00216

Matsu-Ura, T., Shirakawa, H., Suzuki, K. G. N., Miyamoto, A., Sugiura, K., Michikawa, T., Kusumi, A., & Mikoshiba, K. (2019). Dual-FRET imaging of IP(3) and Ca(2+) revealed Ca(2+)-induced IP(3) production maintains long lasting Ca(2+) oscillations in fertilized mouse eggs. Sci Rep, 9(1), 4829. https://doi.org/10.1038/s41598-019-40931-w

Metcalfe, D. D., Baram, D., & Mekori, Y. A. (1997). Mast cells. Physiol Rev, 77(4), 1033–1079. https://doi.org/10.1152/physrev.1997.77.4.1033

Metzger, H., Goetze, A., Kanellopoulos, J., Holowka, D., & Fewtrell, C. (1982). Structure of the high-affinity mast cell receptor for IgE. Fed Proc, 41(1), 8–11.

Mohr, F. C., & Fewtrell, C. (1987). Depolarization of rat basophilic leukemia cells inhibits calcium uptake and exocytosis. J Cell Biol, 104(3), 783–792. https://doi.org/10.1083/jcb.104.3.783

Mukerjee, P., Mysels, K.J.,. (1971). Critical Micellle Concentrations of Aqueous Surfacant Systems.

Narita, S., Goldblum, R. M., Watson, C. S., Brooks, E. G., Estes, D. M., Curran, E. M., & Midoro-Horiuti, T. (2007). Environmental estrogens induce mast cell degranulation and enhance IgE-mediated release of allergic mediators. Environ Health Perspect, 115(1), 48–52. https://doi.org/10.1289/ehp.9378

Nelson, D. L. a. C., M.M. (2017). Lehninger Principles of Biochemistry (7 ed.). W.H. Freeman.

Paddock, B. E., Striegel, A. R., Hui, E., Chapman, E. R., & Reist, N. E. (2008). Ca2+-dependent, phospholipid-binding residues of synaptotagmin are critical for excitation-secretion coupling in vivo. J Neurosci, 28(30), 7458-7466. https://doi.org/10.1523/jneurosci.0197-08.2008

Park, C. J., Song, S. H., Kim, D. H., & Gye, M. C. (2016). Developmental and acute toxicity of cetylpyridinium chloride in Bombina orientalis (Amphibia: Anura). Aquat Toxicol, 177, 446–453. https://doi.org/10.1016/j.aquatox.2016.06.022

Popkin, D. L., Zilka, S., Dimaano, M., Fujioka, H., Rackley, C., Salata, R., Griffith, A., Mukherjee, P. K., Ghannoum, M. A., & Esper, F. (2017). Cetylpyridinium Chloride (CPC) Exhibits Potent, Rapid Activity Against Influenza Viruses in vitro and in vivo. Pathog Immun, 2(2), 252–269. https://doi.org/10.20411/pai.v2i2.200

Pottel, J., Armstrong, D., Zou, L., Fekete, A., Huang, X. P., Torosyan, H., Bednarczyk, D., Whitebread, S., Bhhatarai, B., Liang, G., Jin, H., Ghaemi, S. N., Slocum, S., Lukacs, K. V., Irwin, J. J., Berg, E. L., Giacomini, K. M., Roth, B. L., Shoichet, B. K., & Urban, L. (2020). The activities of drug inactive ingredients on biological targets. Science, 369(6502), 403–413. https://doi.org/10.1126/science.aaz9906

Pubchem. (2023). Cetylpyridinium. https://pubchem.ncbi.nlm.nih.gov/compound/31239

Putney, J. W., Jr. (1986). A model for receptor-regulated calcium entry. Cell Calcium, 7(1), 1–12. https://doi.org/10.1016/0143-4160(86)90026-6

Putney, J. W., Jr. (1990). Capacitative calcium entry revisited. Cell Calcium, 11(10), 611–624. https://doi.org/10.1016/0143-4160(90)90016-n

Raut, P., Sasha R. Weller, Bright Obeng, Brandy L. Soos, Bailey E. West, Christian M. Potts, Suraj Sangroula, Marissa S. Kinney, John E. Burnell, Benjamin L. King, Julie, A. Gosse, & Hess, S. T. (2022). Cetylpyridinium chloride (CPC) reduces zebrafish mortality from influenza infection: Super-resolution microscopy reveals CPC interference with multiple protein interactions with phosphatidylinositol 4,5-bisphosphate in immune function. Toxicology and Applied Pharmacology, 440. https://doi.org/10.1016/j.taap.2022.115913

Ravetch, J. V., & Kinet, J. P. (1991). Fc receptors. Annu Rev Immunol, 9, 457–492. https://doi.org/10.1146/annurev.iy.09.040191.002325

Rawlinson, A., Pollington, S., Walsh, T. F., Lamb, D. J., Marlow, I., Haywood, J., & Wright, P. (2008). Efficacy of two alcohol-free cetylpyridinium chloride mouthwashes - a randomized double-blind crossover study. J Clin Periodontol, 35(3), 230–235. https://doi.org/10.1111/j.1600-051X.2007.01187.x

Roos, J., DiGregorio, P. J., Yeromin, A. V., Ohlsen, K., Lioudyno, M., Zhang, S., Safrina, O., Kozak, J. A., Wagner, S. L., Cahalan, M. D., Veliçelebi, G., & Stauderman, K. A. (2005). STIM1, an essential and conserved component of store-operated Ca2+ channel function. J Cell Biol, 169(3), 435–445. https://doi.org/10.1083/jcb.200502019

Rusan, N. M., Fagerstrom, C. J., Yvon, A. M., & Wadsworth, P. (2001). Cell cycle-dependent changes in microtubule dynamics in living cells expressing green fluorescent protein-alpha tubulin. Mol Biol Cell, 12(4), 971–980. https://doi.org/10.1091/mbc.12.4.971

Sánchez Barrueco, Á., Mateos-Moreno, M. V., Martínez-Beneyto, Y., García-Vázquez, E., Campos González, A., Zapardiel Ferrero, J., Bogoya Castaño, A., Alcalá Rueda, I., Villacampa Aubá, J. M., Cenjor Español, C., Moreno-Parrado, L., Ausina-Márquez, V., García-Esteban, S., Artacho, A., López-Labrador, F. X., Mira, A., & Ferrer, M. D. Effect of oral antiseptics in reducing SARS-CoV-2 infectivity: evidence from a randomized double-blind clinical trial. Emerging Microbes & Infections, 11(1), 1833–1842. https://doi.org/10.1080/22221751.2022.2098059

Sangroula, S., Baez Vasquez, A. Y., Raut, P., Obeng, B., Shim, J. K., Bagley, G. D., West, B. E., Burnell, J. E., Kinney, M. S., Potts, C. M., Weller, S. R., Kelley, J. B., Hess, S. T., & Gosse, J. A. (2020). Triclosan disrupts immune cell function by depressing Ca(2+) influx following acidification of the cytoplasm. Toxicol Appl Pharmacol, 405, 115205. https://doi.org/10.1016/j.taap.2020.115205

Scharenberg, A. M., Humphries, L. A., & Rawlings, D. J. (2007). Calcium signalling and cell-fate choice in B cells. Nat Rev Immunol, 7(10), 778–789. https://doi.org/10.1038/nri2172

Seldin, D. C., Adelman, S., Austen, K. F., Stevens, R. L., Hein, A., Caulfield, J. P., & Woodbury, R. G. (1985). Homology of the rat basophilic leukemia cell and the rat mucosal mast cell. Proc Natl Acad Sci U S A, 82(11), 3871–3875. https://doi.org/10.1073/pnas.82.11.3871

Shi, Y., Luo, H. Q., & Li, N. B. (2011). Determination of the critical premicelle concentration, first critical micelle concentration and second critical micelle concentration of surfactants by resonance Rayleigh scattering method without any probe. Spectrochim Acta A Mol Biomol Spectrosc, 78(5), 1403–1407. https://doi.org/10.1016/j.saa.2011.01.018

Smith-Garvin, J. E., Koretzky, G. A., & Jordan, M. S. (2009). T cell activation. Annu Rev Immunol, 27, 591–619. https://doi.org/10.1146/annurev.immunol.021908.132706

Smith, A. J., Pfeiffer, J. R., Zhang, J., Martinez, A. M., Griffiths, G. M., & Wilson, B. S. (2003). Microtubule-dependent transport of secretory vesicles in RBL-2H3 cells. Traffic, 4(5), 302–312. https://doi.org/10.1034/j.1600-0854.2003.00084.x

Society of Toxicology. (2023). The Toxicologist, supplement to Toxicological Sciences (Vol. 192). Oxford University Press.

Stevens, R. L., & Austen, K. F. (1989). Recent advances in the cellular and molecular biology of mast cells. Immunol Today, 10(11), 381–386. https://doi.org/10.1016/0167-5699(89)90272-7

Sulimenko, V., Hájková, Z., Černohorská, M., Sulimenko, T., Sládková, V., Dráberová, L., Vinopal, S., Dráberová, E., & Dráber, P. (2015). Microtubule nucleation in mouse bone marrow-derived mast cells is regulated by the concerted action of GIT1/βPIX proteins and calcium. J Immunol, 194(9), 4099–4111. https://doi.org/10.4049/jimmunol.1402459

Suzuki, J., Kanemaru, K., Ishii, K., Ohkura, M., Okubo, Y., & Iino, M. (2014). Imaging intraorganellar Ca2+ at subcellular resolution using CEPIA. Nat Commun, 5, 4153. https://doi.org/10.1038/ncomms5153

Swindle, E. J., Metcalfe, D. D., & Coleman, J. W. (2004). Rodent and human mast cells produce functionally significant intracellular reactive oxygen species but not nitric oxide. J Biol Chem, 279(47), 48751–48759. https://doi.org/10.1074/jbc.M409738200

Tadokoro, S., Inoh, Y., Nakanishi, M., & Hirashima, N. (2015). Effects of PIP2 on membrane fusion between mast cell SNARE liposomes mediated by synaptotagmin 2. Biochim Biophys Acta, 1848(10 Pt A), 2290-2294. https://doi.org/10.1016/j.bbamem.2015.06.016

Takekawa, M., Furuno, T., Hirashima, N., & Nakanishi, M. (2012). Mitochondria take up Ca2+ in two steps dependently on store-operated Ca2+ entry in mast cells. Biol Pharm Bull, 35(8), 1354–1360. https://doi.org/10.1248/bpb.b110576

Tasaka, K., Mio, M., Fujisawa, K., & Aoki, I. (1991). Role of microtubules on Ca2+ release from the endoplasmic reticulum and associated histamine release from rat peritoneal mast cells. Biochem Pharmacol, 41(6-7), 1031–1037. https://doi.org/10.1016/0006-2952(91)90211-m

Teng, F., He, T., Huang, S., Bo, C. P., Li, Z., Chang, J. L., Liu, J. Q., Charbonneau, D., Xu, J., Li, R., & Ling, J. Q. (2016). Cetylpyridinium chloride mouth rinses alleviate experimental gingivitis by inhibiting dental plaque maturation. Int J Oral Sci, 8(3), 182–190. https://doi.org/10.1038/ijos.2016.18

Theoharides, T. C., Alysandratos, K. D., Angelidou, A., Delivanis, D. A., Sismanopoulos, N., Zhang, B., Asadi, S., Vasiadi, M., Weng, Z., Miniati, A., & Kalogeromitros, D. (2012). Mast cells and inflammation. Biochim Biophys Acta, 1822(1), 21–33. https://doi.org/10.1016/j.bbadis.2010.12.014

Theoharides, T. C., Stewart, J. M., Panagiotidou, S., & Melamed, I. (2016). Mast cells, brain inflammation and autism. Eur J Pharmacol, 778, 96–102. https://doi.org/10.1016/j.ejphar.2015.03.086

Thrasher, S. M., Scalfone, L. K., Holowka, D., & Appleton, J. A. (2013). In vitro modelling of rat mucosal mast cell function in Trichinella spiralis infection. Parasite Immunol, 35(1), 21–31. https://doi.org/10.1111/pim.12014

Trebak, M., & Kinet, J. P. (2019). Calcium signalling in T cells. Nat Rev Immunol, 19(3), 154–169. https://doi.org/10.1038/s41577-018-0110-7

Tsukioka, M., Iino, M., & Endo, M. (1994). pH dependence of inositol 1,4,5-trisphosphate-induced Ca2+ release in permeabilized smooth muscle cells of the guinea-pig. J Physiol, 475(3), 369-375. https://doi.org/10.1113/jphysiol.1994.sp020078

Urata, C., & Siraganian, R. P. (1985). Pharmacologic modulation of the IgE or Ca2+ ionophore A23187 mediated Ca2+ influx, phospholipase activation, and histamine release in rat basophilic leukemia cells. Int Arch Allergy Appl Immunol, 78(1), 92–100. https://doi.org/10.1159/000233869

Vig, M., Beck, A., Billingsley, J. M., Lis, A., Parvez, S., Peinelt, C., Koomoa, D. L., Soboloff, J., Gill, D. L., Fleig, A., Kinet, J. P., & Penner, R. (2006). CRACM1 multimers form the ion-selective pore of the CRAC channel. Curr Biol, 16(20), 2073–2079. https://doi.org/10.1016/j.cub.2006.08.085

Weatherly, L. M., Kennedy, R. H., Shim, J., & Gosse, J. A. (2013). A microplate assay to assess chemical effects on RBL-2H3 mast cell degranulation: effects of triclosan without use of an organic solvent. J Vis Exp(81), e50671. https://doi.org/10.3791/50671

Weatherly, L. M., Nelson, A. J., Shim, J., Riitano, A. M., Gerson, E. D., Hart, A. J., de Juan-Sanz, J., Ryan, T. A., Sher, R., Hess, S. T., & Gosse, J. A. (2018). Antimicrobial agent triclosan disrupts mitochondrial structure, revealed by super-resolution microscopy, and inhibits mast cell signaling via calcium modulation. Toxicol Appl Pharmacol, 349, 39–54. https://doi.org/10.1016/j.taap.2018.04.005

Weatherly, L. M., Shim, J., Hashmi, H. N., Kennedy, R. H., Hess, S. T., & Gosse, J. A. (2016). Antimicrobial agent triclosan is a proton ionophore uncoupler of mitochondria in living rat and human mast cells and in primary human keratinocytes. J Appl Toxicol, 36(6), 777–789. https://doi.org/10.1002/jat.3209

Weller, R. S., John E. Burnell, Brandon M. Aho, Bright Obeng, Emily L. Ledue, Juyoung K. Shim, Samuel T. Hess, & Gosse, a. J. A. (2022). Mechanisms of Antimicrobial Agent Cetylpyridinium Chloride Mitochondrial Toxicity in Rodent and Primary Human Cells: Super-resolution Microscopy Reveals Nanostructural Disruption. bioRxiv. https://www.biorxiv.org/content/10.1101/2022.09.27.509813v1

Wolf, K. j. (2016). Safety and Effectiveness of Consumer Antiseptics; Topical Antimicrobial Drug Products for Over-the-Counter Human Use. Final rule. Fed Regist, 81(172), 61106–61130.

Xue, L., Li, X., Chen, Q., He, J., Dong, Y., Wang, J., Shen, S., Jia, R., Zang, Q. J., Zhang, T., Li, M., & Geng, Y. (2018). Associations between D3R expression in synovial mast cells and disease activity and oxidant status in patients with rheumatoid arthritis. Clin Rheumatol, 37(10), 2621–2632. https://doi.org/10.1007/s10067-018-4168-1

Yaegaki, K., & Sanada, K. (1992). Biochemical and clinical factors influencing oral malodor in periodontal patients. J Periodontol, 63(9), 783–789. https://doi.org/10.1902/jop.1992.63.9.783

Zaitsu, M., Narita, S., Lambert, K. C., Grady, J. J., Estes, D. M., Curran, E. M., Brooks, E. G., Watson, C. S., Goldblum, R. M., & Midoro-Horiuti, T. (2007). Estradiol activates mast cells via a non-genomic estrogen receptor-alpha and calcium influx. Mol Immunol, 44(8), 1977–1985. https://doi.org/10.1016/j.molimm.2006.09.030

