## Supplement for "Pharmaceutical agent cetylpyridinium chloride inhibits immune mast cell function by interfering with calcium mobilization"

**Table of contents**

**CPC effects on organic voltage-sensitive dye DiSC_3_(5) ……………………… S-1**

**CPC effects on fluorophore used in *in vitro* tubulin assay……………………S-2**

**CPC effects on organic voltage-sensitive dye DiSC_3_(5)**

A no-cell experiment was performed to determine the effect of CPC on the fluorescence of the voltage-sensitive dye 3,3’-dipropylthiadicarbocyanine iodide (DiSC_3_(5)).

**Method:** DiSC_3_(5) (TCI America, 98% purity, CAS no. 53213-94-8) was dissolved in dimethyl sulfoxide (DMSO) (Sigma-Aldrich) and was combined with Tyrode’s buffer to form an aqueous solution of 2.04 μM DiSC_3_(5)/0.02% DMSO, which was plated at 160 μL per well in the experimental wells of a Grenier 96-well black-bottom plate. All plating in this experiment was performed at 37 °C. For background measurements, a solution of 0.02% DMSO in Tyrode’s buffer was plated at 160 μL per well. CPC was dissolved in Tyrode’s buffer, and 160 μL 2x CPC solution per well was plated into the corresponding experimental wells containing the DiSC_3_(5) solution. One set of background wells was treated with 0 μM CPC and the other with 15 μM CPC. The plate was incubated for 10 min at 37 °C before fluorescence was measured in a Synergy 2 (Biotek) microplate reader with 530 ± 20 nm excitation and 645 ± 15 nm emission. Raw fluorescence values (with the 0 µM CPC background subtracted) are plotted against the CPC dose.

**Results:** CPC enhances the fluorescence of the voltage-sensitive dye in a dose-dependent manner (Figure S1) following 10 min incubation of DiSC_3_(5) with CPC. Statistically significant results, compared to 0 μM CPC, are represented by ***p < 0.001, determined by one-way ANOVA followed by Tukey’s post-hoc test.

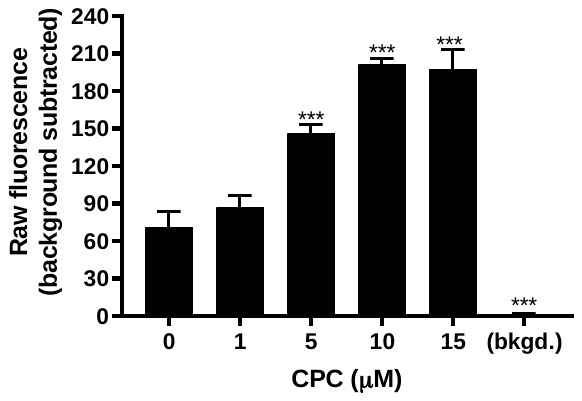

**Figure S1.** **Effects of CPC on DiSC_3_(5) fluorescence.**

Representative data of a no-cell DiSC_3_(5) CPC fluorescence interference assay. Values presented are means ± SD for triplicate measurements in a single representative experiment. Statistically significant results, compared to 0 μM CPC, are represented by ***p < 0.001, determined by one-way ANOVA followed by Tukey’s post-hoc test.

In conclusion, CPC enhances the fluorescence of DiSC_3_(5). As such, DiSC_3_(5) cannot be used as a fluorescent reporter in experiments because CPC modulates its fluorescence.

S-1

**CPC effects on fluorophore used in *in vitro* tubulin assay.**

This control assay aimed to test if CPC, 5 μM, affects the fluorophore used to measure tubulin polymerization.

**Method:** *In vitro* tubulin assay fluorescence was measured per the manufacturer’s instructions as in Methods. The average fluorescence over 60 minutes was measured for this control assay. Each group contains all reagents except for tubulin protein itself. No background subtraction was performed. Data from two independent experiments, one observed in duplicate and a second in triplicate, were averaged. Mean fluorescence intensity was used to compare the control buffer group to the 5 μM CPC group using a two-tailed t-test for the difference between means.
**Results:** CPC, 5 μM, did not interfere with the fluorescence of the dye employed within this kit (Figure S2); p-value > 0.005, determined by a two-tailed t-test for the difference between means (Figure S2).

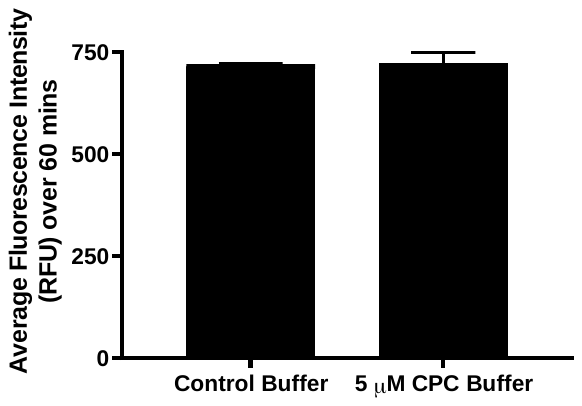

**Figure S2.** **CPC effect on fluorophore used in *in vitro* tubulin polymerization assay**. Fluorescence intensity (y-axis) was measured for one hour to compare the control and 5 µM CPC buffer. Both groups contain active fluorophores but no tubulin protein for polymerization. Data represents average fluorescence intensity ± SD of two independent experiments, one observed induplicate and another in triplicate.

In conclusion, CPC does not interfere with the fluorophore used in the *in vitro* tubulin assay.

S-2
